# Gut Commensals Regulate the Intestinal Kynurenine Pathway

**DOI:** 10.64898/2026.05.08.723850

**Authors:** Ana Djukovic, Chen Liao, Andras Bimbo-Szuhai, Chloe Lindberg, Hui Liu, Hannah Lees, Ruben J. F. Ramos, Justin R. Cross, Joao B. Xavier

## Abstract

The kynurenine pathway (KP) of tryptophan metabolism shapes host immunity and is implicated in inflammatory, malignant, and neurodegenerative diseases. Although it is well known that inflammation influences KP activity, whether and how gut microbiota regulates KP in the intestine remains unclear. Here, we find that in allogeneic hematopoietic cell transplant patients, the kynurenine-to-tryptophan ratio tracks plasma neopterin, whereas intestinal levels do not, suggesting that local KP regulation is not explained by inflammation alone. In mice, targeted microbiota perturbation downregulates intestinal expression of indoleamine 2,3-dioxygenase 1 (IDO1), the rate-limiting enzyme of the KP, while microbiota restoration reversed this effect. Computational analyses associate Lachnospiraceae and Oscillospiraceae with colonic IDO1 in mice and inflammatory bowel disease cohorts. *In vitro* experiments support a mechanism whereby microbiota perturbation alters host absorption of the vitamin E isoform, γ-tocotrienol, which impacts IDO1 expression. Our findings support that the gut microbiota controls colonic KP through micronutrient absorption.

## INTRODUCTION

The composition of the gut microbiota influences host metabolism^1–6^ and immune function^7–10^. Microbiota perturbations, caused by antibiotic exposure or dietary changes, can impair immune homeostasis and contribute to chronic metabolic^11,12^ and inflammatory disorders^13,14^, as well as delay recovery in patients receiving immunosuppressive therapies^10^. Despite extensive evidence of the microbiota-metabolism-immunity axis, the underlying mechanisms linking these processes remain incompletely understood. Addressing this knowledge gap can lead to the development of novel microbiota-based interventions to modulate metabolism and control immune responses.

The kynurenine pathway (KP) is the principal route of tryptophan catabolism and has special clinical relevance due to its role in immunity^15–20^. Its activity is stimulated by pro-inflammatory cytokines^21^. Moreover, kynurenine, the first intermediate in the pathway, suppresses the proliferation and survival of effector T cells^22^ and acts as a ligand for the aryl hydrocarbon receptor (AhR), promoting the differentiation of naïve CD4+ T cells into Foxp3+ regulatory T cells^23^, important for immune tolerance. Other KP products, such as 3-hydroxykynurenine, also exhibit immunosuppressive properties^24^, while others play key physiological roles: quinolinic acid acts as an N-methyl-D-aspartate (NMDA) receptor agonist, and nicotinamide adenine dinucleotide (NAD^+^) is an essential cellular cofactor^25^. This means that perturbations of the KP—through the microbiota or other means—can have broad consequences in health and disease.

The conversion of tryptophan to kynurenine, the first and rate-limiting step in KP, is catalyzed by three enzymes: tryptophan 2,3-dioxygenase (TDO), constitutively expressed in the liver, and by indoleamine-2,3-dioxygenases 1 and 2 (IDO1 and IDO2), which are expressed in extrahepatic tissues^25^. Of these, IDO1 is inducible by pro-inflammatory cytokines^21^ and highly expressed in immune cells and epithelial tissues, including the intestinal mucosa. Therefore, colonic expression of IDO1 could be a key determinant of the KP flux in the gut. Emerging evidence suggests that the gut microbiota can modulate KP activity. Germ-free mice exhibit a reduced kynurenine-to-tryptophan (KYN/TRP) ratio compared to conventionally colonized mice^26^. Additionally, administration of probiotic strains has been reported to either upregulate or downregulate IDO1 expression, depending on the bacterial species, and on the host and its health status^27–29^. However, the role of commensal strains in KP activity and how they might regulate it are still largely unclear.

In this study, we integrate patient-derived datasets, mouse models, computational analyses, metabolomics, and *in vitro* mechanistic approaches to investigate the role of commensal gut microbes on intestinal IDO1 expression and activity of tryptophan catabolism through the KP. We identify members of the Lachnospiraceae and Oscillospiraceae families as key microbial drivers of colonic IDO1 expression and uncover a microbiome-dependent mechanism in which absorption of γ-tocotrienol (γT3), a vitamin E isoform, is modulated by gut bacteria to impact IDO1 expression.

Together, these findings reveal a previously unrecognized axis linking gut commensals, micronutrient bioavailability, and host tryptophan catabolism, a link with broad implications for host physiology, including diseases such as inflammatory disorders and cancer.

## RESULTS

### Patient analyses suggest that KP activity in the intestine is independent from systemic inflammation

We recently published an atlas of fecal microbiome compositions from over 2,000 patients who underwent allogeneic hematopoietic cell transplantation (allo-HCT) at Memorial Sloan Kettering Cancer Center^30^. From this resource, we retrospectively selected 92 stool samples representing both biodiverse microbiota states and antibiotic-perturbed states, characterized by low biodiversity and domination by a single taxon (**Figure 1A**). Then, we used a targeted LC-MS/MS panel to quantify tryptophan (TRP), kynurenine (KYN), and neopterin (a marker of immune activation) in paired fecal and plasma samples. Consistent with the known induction of the KP by proinflammatory cytokines^21^, the neopterin levels in patients’ plasma were positively correlated with their systemic KYN/TRP ratio (**Figure 1B**, ρ=0.60, P = 10^-5^). In contrast, no positive correlation was observed between stool neopterin and gut KYN/TRP. In fact, the correlation was mildly negative (**Figure 1C**, ρ=-0.22, P < 0.05), suggesting that the gut microbiota may impact the activity of the KP in the intestine.

**Figure 1.**
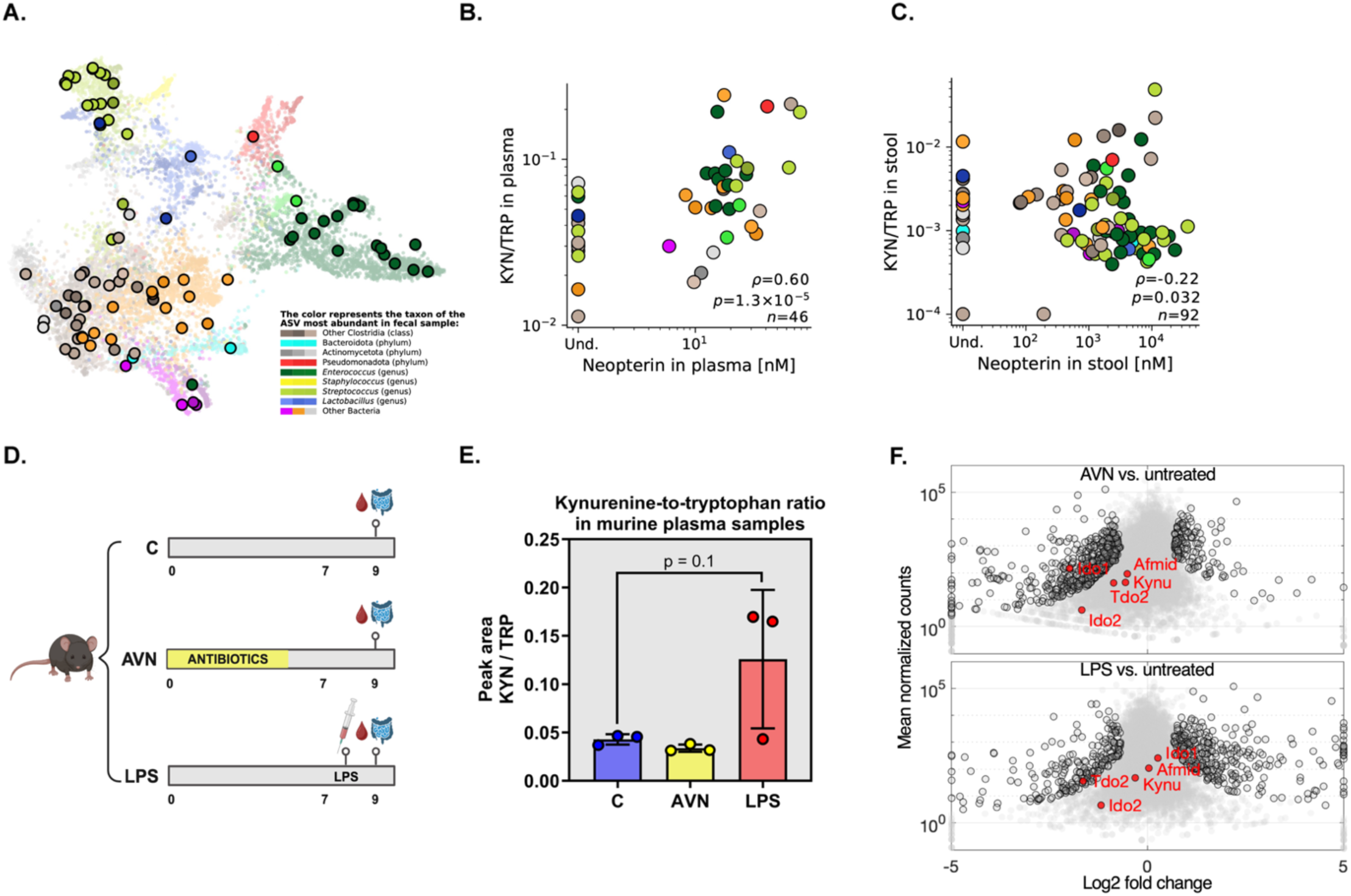
Systemic kynurenine pathway is inflammation driven, while local gut flux depends on the microbiome. A) TaxUMAP embedding of the allo-HCT fecal microbiome atlas, with the 92 stool samples selected for paired stool and plasma metabolite profiling highlighted with black outlines. Point colors indicate the taxon of the dominant amplicon sequence variant (ASV) in each sample. B) Plasma neopterin versus plasma KYN/TRP ratio for the subset of 92 samples, sowing a positive association between systemic immune activation and systemic KP activity (Spearman ρ = 0.60, P = 0.032). C) Stool neopterin versus stool KYN/TRP ratio across the selected cohort, showing no positive association between immune activation and local KP activity (Spearman ρ = -0.22, P = 1.29 x 10^-5^). D) Schematic representation of the experiment in which antibiotic cocktail and LPS were given to mice to disentangle inflammation- from microbiome-mediated KP regulation. AVN: ampicillin, vancomycin, and neomycin; LPS: lipopolysaccharide. E) LPS-induced inflammation increases systemic KYN/TRP ratio, while antibiotics-mediated microbiome perturbation (AVN) does not. P = 0.1, one-way ANOVA. F) RNA sequencing of the small intestine and gene enrichment analysis show that local KP is regulated by the gut microbiota. Antibiotic-cocktail treated mice (AVN) showed significant downregulation of the local KP (GO:0019441). Adjusted P < 0.01, hypergeometric probability density function. On the other hand, LPS treatment had no effect on local KP (GO:0019441). Adjusted P > 0.05, hypergeometric probability density function. Each dot represents one gene transcript. Negative log2 fold change between mean normalized counts of untreated and antibiotic-cocktail/LPS treated animals reveals down-regulation. Transcripts with outline were significantly different between experimental groups (adjusted P > 0.05, negative binomial test). Transcripts of the genes from the KP (GO:0019441) are marked in red. In bar plot, bars represent mean, and error bars represent the standard deviation.

### Experiments in mice link shifts in the gut microbiota to KP in the intestine, independently of immune activation

Allo-HCT patients experience extreme immune profile changes and gut microbiota shifts simultaneously^10^, making it impossible to distinguish whether KP regulation is immune- or microbiome-driven. Therefore, to address this limitation, we used a controlled mouse experiment where we perturbed the microbiota or the immune system independently (**Figure 1D**). One group of animals received intraperitoneal lipopolysaccharide (LPS) to induce systemic inflammation, while another received a cocktail of ampicillin, vancomycin and neomycin (AVN) in drinking water for one week–a regimen that markedly depletes the gut microbiota^31,32^. Plasma metabolic profiling revealed increased plasma KYN/TRP in two out of three mice that received systemic immune activation through LPS treatment, consistent with inflammation-driven KP activation via IDO1 (**Figure 1E**, P = 0.1, one-way ANOVA). In contrast, AVN treatment had no effect on the systemic KYN/TRP (**Figure 1E**, P > 0.05, one-way ANOVA). RNA sequencing of intestinal tissue and gene set enrichment analysis revealed that AVN treatment downregulated the KP (GO:0019441, **Figure 1F**, adjusted P < 0.01, hypergeometric probability density function), lowering expression of AFMID, KYNU, TDO2, IDO1, and IDO2, with IDO1 showing the most pronounced reduction (**Figure 1F**, adjusted P < 0.001, negative binomial test). However, the LPS treatment had no effect on the local KP (GO:0019441, **Figure 1F**, adjusted P > 0.05, hypergeometric probability density function).

These results indicated that systemic KP activity can be driven by immune activation, while the intestinal KP activity appears to be dependent on the gut microbiome perturbations, and not systemic inflammation.

### Microbiota perturbation downregulates IDO1 in colonocytes

We next focused on measuring expression of the IDO1, responsible for the rate-limiting step of the KP and expressed in intestinal mucosa. Previous meta-analysis has shown that IDO1 expression is correlated with the abundance of kynurenine across human tissues and organs^33^. In fact, this is one of the strongest mRNA-metabolite correlations observed in humans, making IDO1 a reliable proxy for local KP activity, and a valuable readout for this pathway’s activity. Given the strong reduction of IDO1 expression in AVN-treated mice, we first examined whether this effect was driven by a migration of IDO1-expressing cells (e.g., immune cells) away from the colon or by a downregulation of IDO1 expression in resident intestinal cells.

We again treated C57BL/6J mice with the AVN cocktail in drinking water for 1 week (**Figure 2A**). 16S rDNA qPCR analysis of fecal samples collected one day post-treatment confirmed a significant reduction in total bacterial load compared with controls (**Figure 2B**, P < 0.0001, Mann-Whitney test, two independent experiments). RT-qPCR analysis of colonic tissue revealed an ∼83% reduction in IDO1 mRNA expression compared with untreated animals (**Figure 2C**, P < 0.0001, Mann-Whitney test, two independent experiments), consistent with our previous RNA-seq results (**Figure 1F**). We then performed fluorescent *in-situ* hybridization (FISH) for IDO1 on colonic cross-sections (**Figure 2D**) and co-stained with E-cadherin to identify epithelial cells. We saw a significantly higher portion of IDO1-positive cells among colonocytes compared to non-epithelial IDO1-positive cells (**Figure 2E**, P < 0.0001, Mann-Whitney test). In both control and antibiotic-treated animals, more than 60% of IDO1-positive cells were colonocytes (**Figure 2E**), indicating that epithelial cells are the predominant source of IDO1 in this model. Antibiotic treatment reduced IDO1 fluorescence intensity without reducing the number of IDO1-positive cells (**Figure S1**, P = 0.55, Mann-Whitney test), but rather through downregulation within existing IDO1-expressing cells (**Figure 2F**, P < 0.05, Mann-Whitney test).

**Figure 2.**
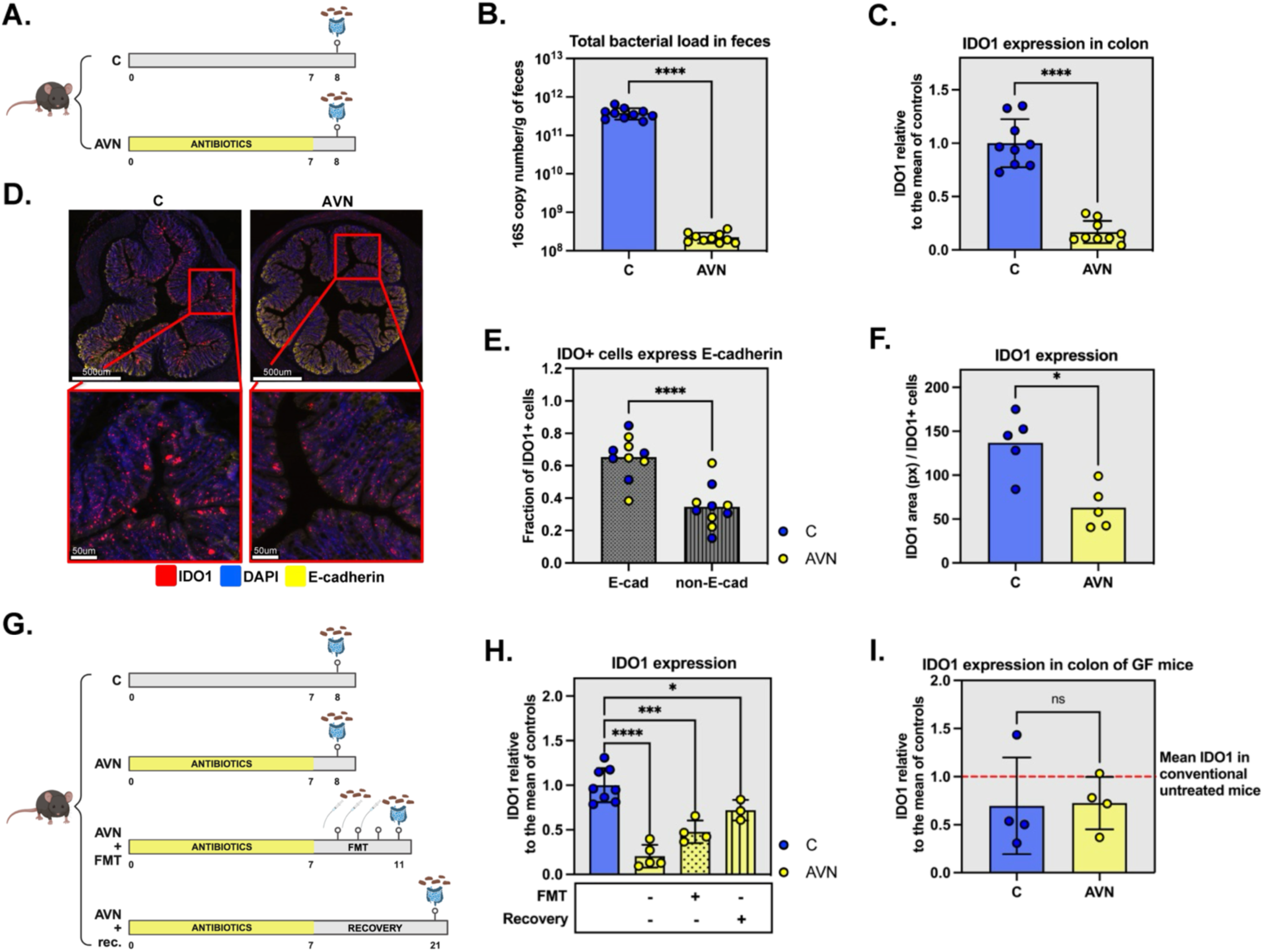
Antibiotic treatment downregulates IDO1 in colonocytes via modulation of the microbiome. A) Schematic representation of the experiment in which antibiotic cocktail was administered to mice to perturb their gut microbiome. AVN: ampicillin, vancomycin, and neomycin. B) The antibiotic cocktail depletes gut bacteria, as reflected by 16S rDNA copy number per gram of feces. C-untreated mice, AVN-antibiotic treated mice. P < 0.0001, two-tailed Mann-Whitney test, two independent experiments. C) The antibiotic cocktail downregulates IDO1 in colonic tissue. IDO1 levels in colonic tissue were normalized by the levels of the endogenous control (HMBS), and the mean IDO1 expression in control samples. P < 0.0001, two-tailed Mann-Whitney test, two independent experiments. D) Fluorescent *in-situ* hybridization of IDO1 in colonic cross-sections from control (C) and antibiotic cocktail-treated animals (AVN). Red: IDO1, blue: DAPI, yellow: E-cadherin. E) IDO1 is primarily expressed in colonocytes. Fraction of IDO1-positive cells that also express E-cadherin (E-cad) versus those that do not (non-E-cad). Blue circles represent samples from untreated animals, while yellow circles represent samples from AVN-treated animals. P < 0.0001, two-tailed Mann-Whitney test. F) Decrease in IDO1 expression levels is due to reduced transcription. The area covered by the IDO1 probe (in pixels) in IDO1-positive cells in untreated (C) and AVN-treated animals. P < 0.05, two-tailed Mann-Whitney test. G) Schematic representation of the experiment in which IDO1 expression was measured after different microbiome-recovery approaches in antibiotic-treated animals. AVN: ampicillin, vancomycin, and neomycin cocktail. FMT: fecal microbiome transplant. H) Microbiome recovery restores IDO1 expression levels. IDO1 levels in colonic tissue were normalized by the levels of the endogenous control (HMBS), and the mean IDO1 expression in control samples. * P < 0.05, *** P < 0.001, **** P < 0.0001, one-way ANOVA. I) The antibiotic cocktail does not affect IDO1 levels in germ-free animals. Red line represents IDO1 expression in conventional, untreated mice. P = 0.69, two-tailed Mann-Whitney test, two independent experiments. In all plots, bars represent mean, and error bars represent the standard deviation. See also **Figure S1**.

These results indicate that immune cells were not the main source of IDO1 in the colon and that the reduction in IDO1 in AVN-treated mice was due to its downregulation, and not migration of IDO1-expressing immune cells away from the colon.

### Restoring commensal populations normalizes colonic IDO1 expression

To confirm the role of the gut microbiota in regulating colonic IDO1, we first treated three groups of mice with the AVN cocktail and then evaluated the impact of microbiota recovery on colonic IDO1 (**Figure 2G**). The first group (the control group) was analyzed the day after the antibiotic cessation, as in the previous experiment. As before, the antibiotics reduced IDO1 expression by approximately 80% (**Figure 2H**, P < 0.0001, one-way ANOVA). The second group (the FMT group) received oral gavage with a fecal slurry from untreated animals for three consecutive days post-antibiotics. The third group (the recovery group) was allowed to recover spontaneously for 2 weeks after antibiotics. FMT partially restored IDO1 by day 10 (P < 0.001, one-way ANOVA), and spontaneous recovery led to an even greater recovery of IDO1 by day 20 (P < 0.05, one-way ANOVA). In summary, relative to the AVN-induced decrease, FMT and recovery restored ∼34% and ∼65% of the lost signal, respectively.

To exclude the possibility of a direct antibiotic effect on IDO1 expression, we treated germ-free mice with the AVN cocktail for 1 week (**Figure 2I**). IDO1 levels in antibiotic-treated germ-free mice were indistinguishable from those of untreated germ-free controls (P = 0.69, Mann-Whitney test, two independent experiments). Taken together, these experiments confirmed that the downregulation observed in conventional mice is microbiota-dependent.

### Computational analysis identifies commensal taxa associated with colonic IDO1 expression

To identify the specific commensal taxa whose antibiotic-mediated perturbation may explain a decrease in colonic IDO1 expression, we treated 54 conventional mice with a range of antibiotic regimens, including single agents (ampicillin, vancomycin, metronidazole, ciprofloxacin), the AVN cocktail, and included the endpoints of the microbiota recovery conditions described above (**Figure 2G**). For each mouse, we quantified IDO1 mRNA in colonic tissue and profiled fecal microbiota using 16S rDNA gene sequencing (**Figure 3A**).

**Figure 3.**
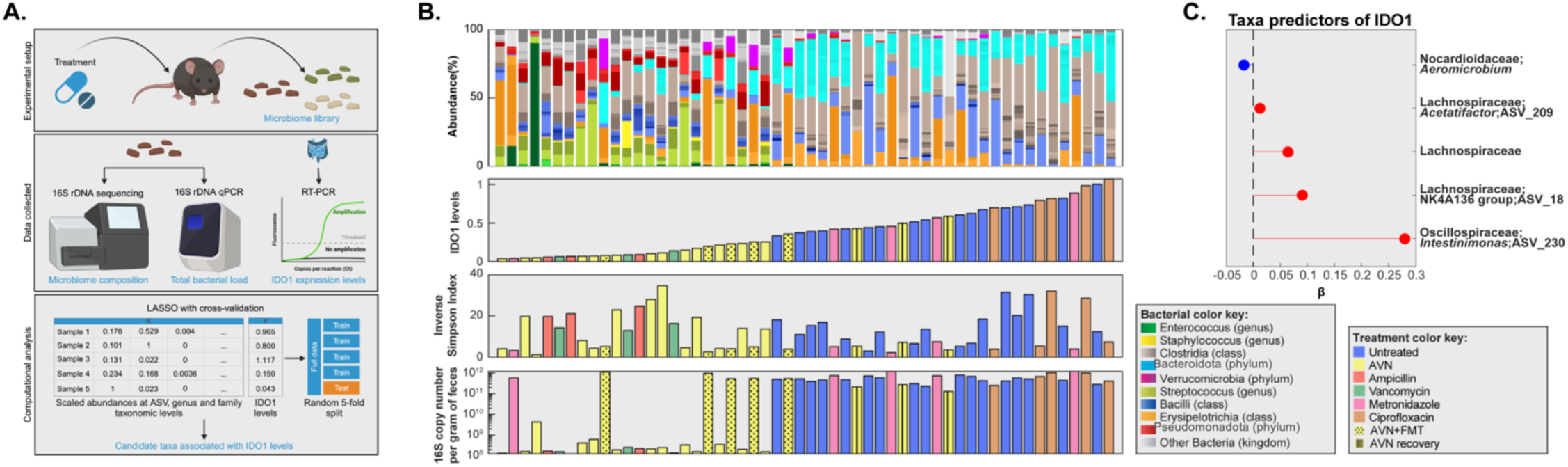
Computational analysis of microbiome perturbation experiments reveals candidate bacterial modulators of the kynurenine pathway. A) Schematic representation of the strategy used to identify microbiome members that correlate with IDO1 expression in colon. B) Distinct microbiome composition patterns correspond to low and high IDO1 expression in the colon, whereas bacterial load does not. Each bar represents one animal. The panels (from top to bottom) show microbiome composition, colonic IDO1 levels, bacterial diversity (measured by the Inverse Simpson Index), and bacterial load (measured by 16S rDNA copy number per gram of feces). Bars are sorted by IDO1 expression levels from lowest to highest. Colors in the top panel represent the relative abundances of different microbiome members, as indicated in the “Bacterial color key”. Colors of the second, third and fourth panels indicate the treatment each animal received, as shown in the “Treatment color key”. C) Taxa selected by LASSO regression as predictors of IDO1 levels in the colon. Red indicates taxa positively correlated with IDO1 levels, while blue represents taxa negatively correlated. β: LASSO coefficient. ASV: amplicon sequence variant. See also **Figure S2**, and **Figure S3.**

These treatments generated a wide spectrum of microbiome compositions (**Figure 3B**, top panel), and their paired IDO1 colonic expressions (**Figure 3B**, second panel from top). The microbiota biodiversity quantified by the Inverse Simpson Index (second panel from bottom), and the total bacterial load (bottom panel) varied widely among them. When samples were sorted by IDO1 expression, it became evident that bacterial diversity was uncorrelated with the colonic IDO1 levels (Pearson correlation, R^2^ = 0.0039, P = 0.65). In contrast, total bacterial load explained ∼50% of the variation in IDO1 expression (Pearson correlation, R^2^ = 0.50, P < 0.0001). Still, there were notable exceptions: some FMT-treated mice had high bacterial loads but only partial IDO1 recovery, suggesting that bacterial load alone is insufficient for normal IDO1 expression. Visual inspection of microbiome profiles suggested that mice with high IDO1 levels tended to harbor more members of the phylum Bacteroidota (shown in cyan) and the class Bacilli (in blue), whereas low-IDO1 mice were enriched in Pseudomonadota (in red) and *Streptococcus* (in lime green).

To identify taxa associated with IDO1 expression, we applied least <u>a</u>bsolute <u>s</u>hrinkage and <u>s</u>election <u>o</u>perator (LASSO) regression using scaled relative abundances at the ASV, genus, and family levels as predictors. The LASSO selected four taxa positively associated with IDO1 and one negatively associated (**Figure 3C**, **Figure S2**). A linear regression model fitted with these taxa explained 79% of IDO1 variability (P < 0.0001, adjusted R^2^ = 0.79). The strongest positive effect was for an ASV classified as *Intestinimonas* (family Oscillospiraceae), followed by ASVs classified as *Acetatifactor* and NK4A136 group (both Lachnospiraceae), and the Lachnospiraceae family. These associations, discovered using bacterial relative abundances, held when using absolute abundances (relative abundance multiplied by total 16S rDNA gene copies), with *Intestinimonas* (ASV230) remaining the strongest predictor (**Figure S3**). Additional predictors in this model included *Intestinimonas*, an ASV classified as *Colidextribacter* (family Oscillospiraceae) and UCG_010 (family Ruminococcaceae). This set of predictors explained 67% of the variability in colonic IDO1 levels when used to fit a linear regression model (P < 0.0001, R^2^ adjusted = 0.67%). Together, these findings show that, at least in mice, IDO1 colonic levels can be predicted from microbiota composition, specifically from the abundances of members of the Lachnospiraceae and Oscillospiraceae families. These commensals are associated with higher IDO1 expression, and their perturbation by antibiotics could therefore lower the KP activity.

### Lachnospiraceae and Oscillospiraceae mediated restoration of colonic IDO1 expression is linked to luminal γ-tocotrienol (γT3) levels

Given that Lachnospiraceae and Oscillospiraceae correlate with IDO1 expression in our mouse model, we next tested whether they could restore IDO1 expression in antibiotic-treated mice. Mice previously treated with AVN cocktail were orally gavaged with a mixture of three Lachnospiraceae and two Oscillospiraceae strains, comprising both human and murine isolates available in our lab and closely related to taxa identified in predictive analyses (**Figure 4A**).

**Figure 4.**
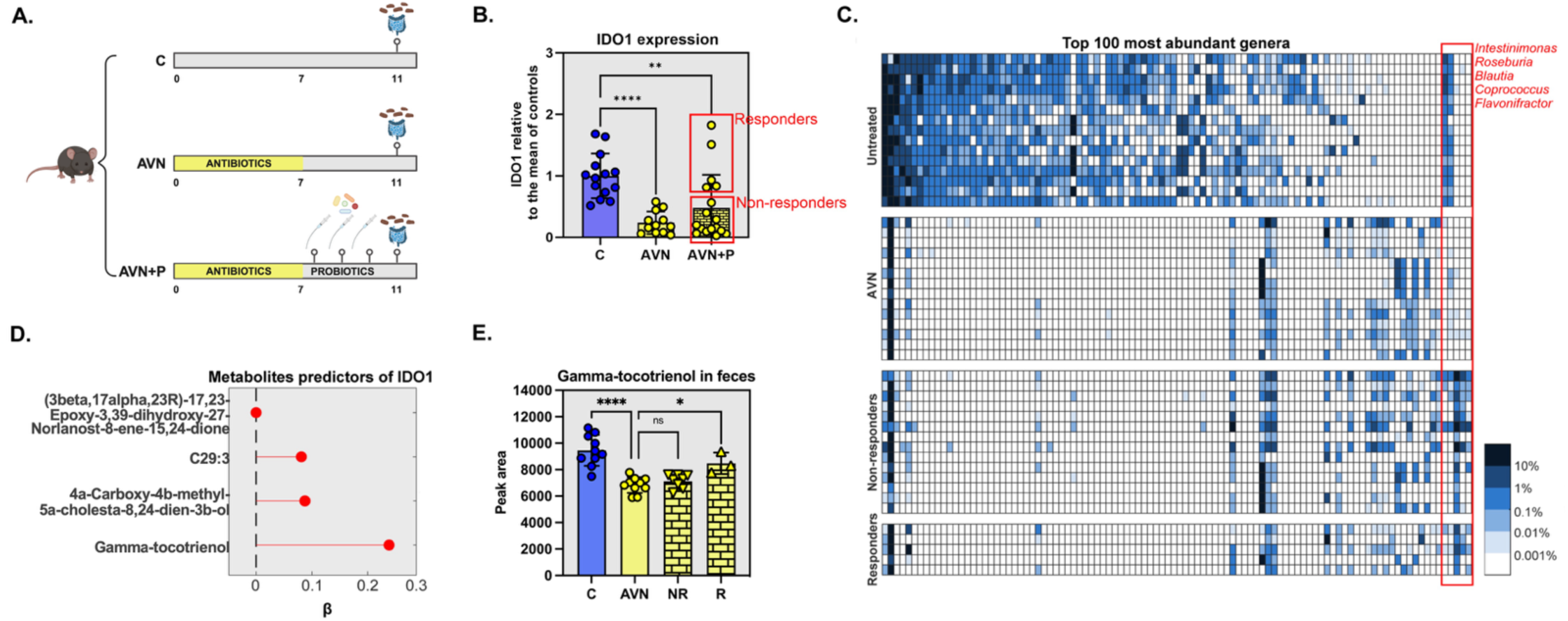
Variable response to probiotic treatment for restoring colonic IDO1 levels is linked to the levels of γ-tocotrienol (γT3), a form of vitamin E. A) Schematic representation of the experiment in which antibiotic cocktail-treated animals received a mix of three Lachnospiraceae and two Oscillospiraceae strains to restore IDO1 levels. AVN: ampicillin, vancomycin, and neomycin cocktail. B) Probiotic treatment restored IDO1 in some, but not all, antibiotic cocktail-treated animals. IDO1 levels in colonic tissue were normalized by the levels of the endogenous control (HMBS), and the mean IDO1 expression in control samples. ** P < 0.01, **** P < 0.0001, one-way ANOVA, three independent experiments. C) Heatmap showing top 100 most abundant genera in fecal samples collected from these experiments. The color bar represents relative abundance cutoffs. Administered strains are marked in red. D) Fecal metabolites selected by LASSO regression as predictors of colonic IDO1 levels. All predictors show a positive correlation with IDO1. β: LASSO coefficient. E) γT3 levels in cecal content decrease after antibiotic cocktail treatment (AVN) and increase in probiotic responders (R), while remaining low in non-responders (NR). Ns: not significant, * P < 0.05, **** P < 0.0001, one-way ANOVA, three independent experiments. In all bar plots, bars represent mean, and error bars represent the standard deviation. See also **Figure S4**, and **Table S1**.

Across three independent experiments, oral administration of these strains recovered colonic IDO1 expression, but only in a subset of the antibiotic-treated mice (**Figure 4B**, three independent experiments). Animals were therefore classified as **responders** (IDO1 restored) or **non-responders** (IDO1 unchanged), with the responder/non-responder pattern reproducible across experiments.

We then asked whether mouse-to-mouse differences in either the colonization efficiency of the administered strains or the recovery of other microbiome members could explain why some mice responded but others not. The gavaged strains, detected by 16S rDNA sequencing, showed similar abundances in both responders and non-responders (**Figure 4C**). There were also no other differences in microbiota composition (**Table S1**, P > 0.05, Wilcoxon test with false discovery rate-FDR), indicating that the difference in response was not due to a variation in bacterial engraftment or remaining microbiome recovery.

Next, we compared the chemical composition of the fecal samples from responders and non-responders. Untargeted metabolomics of cecal contents, followed by LASSO regression (**Figure 4D, Figure S4**), identified four metabolites positively associated with colonic IDO1 expression. The strongest predictor was γ-tocotrienol (γT3), a vitamin E isoform. Antibiotic treatment significantly reduced cecal γT3 levels (**Figure 4E**, P < 0.0001, one-way Anova, two independent experiments), which remained low in non-responders (**Figure 4E**, P > 0.05, one-way Anova, two independent experiments), but recovered in responders (**Figure 4E**, P < 0.05, one-way ANOVA, two independent experiments). Together, these findings suggest that the capacity of Lachnospiraceae and Oscillospiraceae strains to restore IDO1 expression in antibiotic-treated mice is linked not to engraftment success or the recovery of the remaining microbiome, but to the variation in the levels of a micronutrient: γT3.

### Microbiota modulation of γT3 uptake impacts IDO1 expression

To assess the effect of γT3 on IDO1 expression directly in host cells, we established an *in vitro* model using the murine colorectal cancer cell line MC-38. Previous studies have shown that while IDO1 is readily detectable in tumor tissues, its expression is markedly reduced under standard two-dimensional culture conditions, but can be restored by culturing cells in low-attachment plates that promote 3-D spheroid formation^34^. Consistent with this, IDO1 expression was significantly higher in MC-38 cells cultured under low-attachment conditions compared with standard monolayer cultures (**Figure 5A**, P < 0.05, one-tailed Man-Whitney test). We therefore used this 3-D culture system for all subsequent *in vitro* experiments. Next, to confirm that IDO1 expression in this model remained responsive to known regulatory stimuli, MC-38 spheroids were exposed to interferon-gamma (IFNγ)^35^. As expected, these cells, exposed to IFNγ, upregulated IDO1 expression (**Figure 5B**, P < 0.05, one-tailed Man-Whitney test).

**Figure 5.**
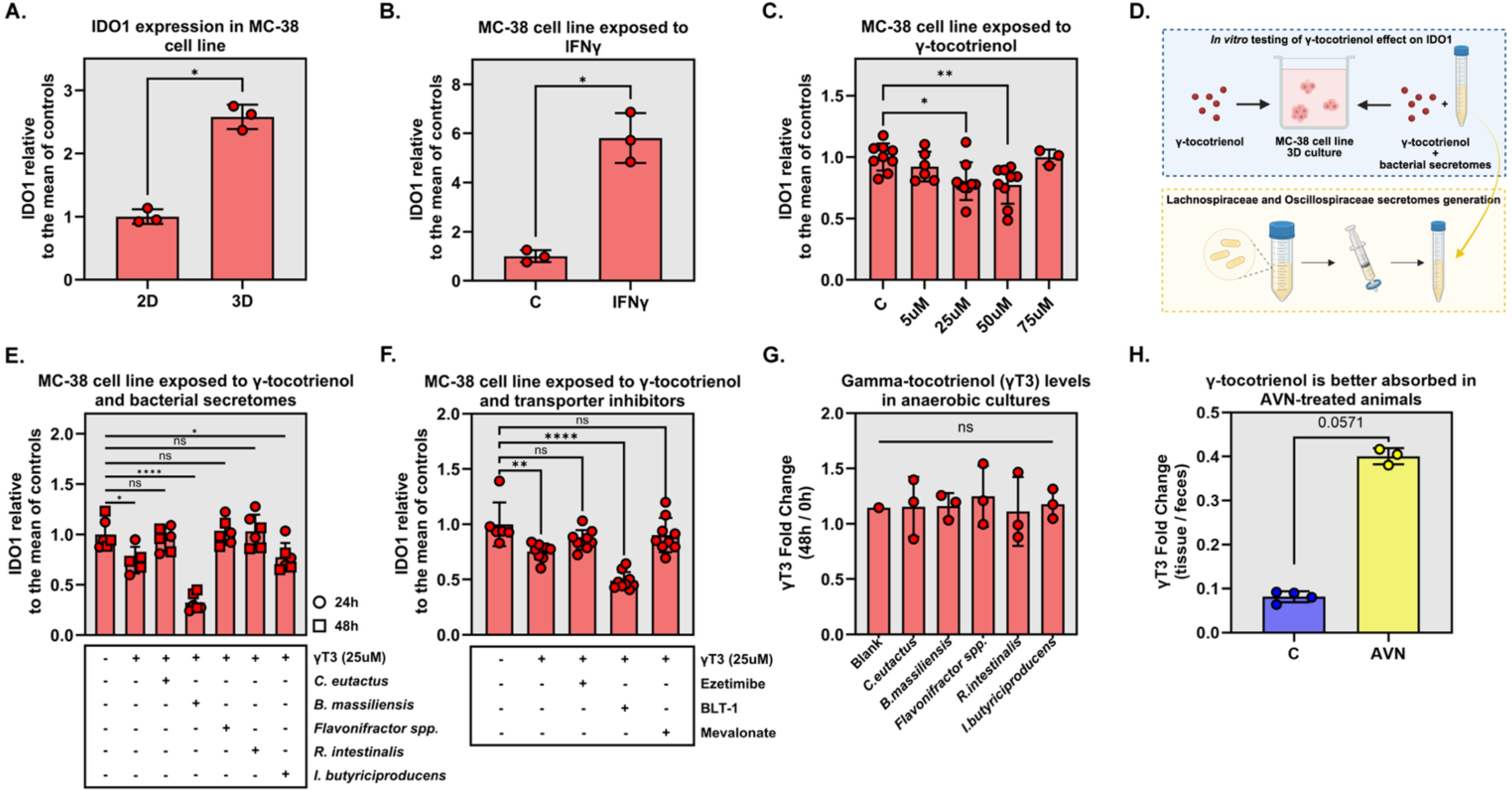
γT3 downregulates IDO1 *in vitro*, and the effect can be reversed by exposure to Lachnospiraceae and Oscillospiraceae secretomes. A) Establishing the model for investigating IDO1 expression *in vitro*. Culturing of MC-38 mouse colorectal cancer cell line in low-attachment plates (3D) led to more robust expression of IDO1 when compared to standard culture conditions (2D). P = 0.05, one-tailed Mann-Whitney test. B) IDO1 expression in MC-38 cell line is inducible by IFNγ. P = 0.05, one-tailed Man-Whitney test. C) Exposure of MC-38 colorectal cancer cell line to γT3 revealed dose-dependent IDO1 downregulation. * P < 0.05, ** P < 0.01, one-way ANOVA, three independent experiments. D) Schematic representation of *in vitro* experiments investigating the effect of γT3, alone or in combination with Lachnospiraceae and Oscillospiraceae secretomes, on MC-38 mouse colorectal cancer cell line. E) Some secretomes restore IDO1 expression in MC-38 cells exposed to γT3. Lachnospiraceae and Oscillospiraceae secretomes were added to MC-38 cells exposed to γT3 for 24 hours or 48 hours periods, revealing that some of the strains’ supernatants can restore IDO1 levels in the presence of γT3. Ns- not significant, * P <0.05, **** P < 0.0001, one-way ANOVA. F) The effect of γT3 on IDO1 expression in MC-38 cell line can be restored by supplementation of transport inhibitor ezetimibe or mevalonate. Ns- not significant, ** P < 0.01, **** P < 0.0001, one-way ANOVA, three independent experiments. G) Lachnospiraceae and Oscillospiraceae strains do not produce nor consume γT3. Five bacterial strains used to restore colonic IDO1 in mice were grown anaerobically in cecal content filtrates spiked with γT3 (for detection purposes). Comparison of γT3 fold changes, calculated by dividing γT3 levels at 48 hours and at the beginning of incubation, revealed no significant differences between γT3 levels in different cultures and blank media. Ns: not significant, P > 0.05, one-way ANOVA. H) AVN-treated animals show higher absorption of γT3. Analysis of γT3 levels in cecal content and tissue of untreated and AVN-treated animals revealed better absorption of γT3 in antibiotic-treated animals, as witnessed by higher fold change between γT3 tissue and cecal content levels. P = 0.057, two-tailed Mann-Whitney test. In all bar plots, bars represent mean, and error bars represent the standard deviation.

Having established a robust model, we next tested the direct effect of γT3 on IDO1 expression (**Figure 5D**). The response to γT3 was non-monotonic, but included a dose-dependent reduction in IDO1 expression in the 5-50μM range (**Figure 5C**, P < 0.05, one-way ANOVA, three independent experiments). This suggests that γT3 has a complex effect, including a potential negative regulatory role on IDO1 expression in colorectal cancer cells.

We then investigated whether secreted metabolites from Lachnospiraceae and Oscillospiraceae strains could modulate γT3-mediated IDO1 downregulation. Bacterial strains were cultured in cell culture medium, supernatants were filtered to remove bacterial cells, and MC-38 spheroids were treated with γT3 in the presence of bacterial secretomes for 24 or 48 hours (**Figure 5D**). Secretomes from *Roseburia intestinalis*, *Coprococcus eutactus*, and *Flavonifractor spp.* reversed the inhibitory effect of γT3 on IDO1 expression, whereas *Blautia massiliensis* further enhanced IDO1 suppression and *Intestinimonas butyriciproducens* showed no effect (**Figure 5E**, P > 0.05, one-way ANOVA). Notably, the effects of *R. intestinalis* and *C. eutactus* were γT3-dependent, as exposure to bacterial supernatants alone did not alter IDO1 expression (**Figure S5**, P > 0.05, one-way ANOVA). These findings demonstrate that secreted microbial products can differentially modulate γT3 activity, revealing an additional layer of microbiota-mediated regulation of host IDO1 expression.

Next, we tested whether inhibition of γT3 uptake reverses its suppressive effect on IDO1 expression. Co-treatment of MC-38 spheroids with γT3 and ezetimibe, an inhibitor of a γT3 transporter, restored IDO1 expression to levels comparable to γT3-untreated controls (**Figure 5F**, P > 0.05, one-way ANOVA, three independent experiments). In contrast, inhibition of an alternative γT3 transporter using BLT-1 potentiated γT3-mediated IDO1 suppression. γT3 has previously been shown to inhibit 3-hydroxy-3-methylglutaryl-coenzyme A (HMG-CoA) reductase, a key enzyme in the mevalonate pathway^36^. Additionally, another study demonstrated that HMG-CoA inhibition downregulates activation of nuclear factor kappa B (NF-kB)^37^, a major driver of IDO1 transcription. To test whether γT3-mediated IDO1 suppression in our model was dependent on HMG-CoA reductase inhibition, we supplemented γT3-treated MC-38 spheroids with mevalonate, the product of HMG-CoA reductase. Mevalonate supplementation completely abrogated the suppressive effect of γT3 on IDO1 expression (**Figure 5F**, P > 0.05, one-way ANOVA, three independent experiments), indicating that γT3 downregulates IDO1 via inhibition of the mevalonate pathway.

Finally, having established γT3 as a modulator of IDO1 expression, we wondered what the link between Lachnospiraceae/Oscillospiraceae strains and γT3 levels in the gut is. Diet is the only known source of γT3^38^. Since all mice in our experiments received identical chow and showed no differences in food intake, variation in diet does not explain the γT3 luminal variability that correlated with IDO1 expression in responders. Therefore, we explored whether the gavaged strains produced γT3 or, instead, modulated its levels by altering absorption. To test direct production or degradation, each strain was cultured anaerobically in filtered cecal content supplemented with a small amount of γT3 (for detection accuracy). After 48 hours, the fold change in γT3 did not differ between different cultures and uninoculated blank media (**Figure 5G**, P > 0.05, one-way ANOVA), indicating that these strains neither synthesize nor degrade γT3.

We next returned to mouse model to investigate whether antibiotic-induced microbiota perturbation impacted γT3 absorption *in vivo*. We compared the γT3 levels in cecal contents and corresponding tissues from AVN-treated and untreated mice. AVN-treated animals exhibited increased γT3 absorption, reflected by a higher tissue-to-content ratio (**Figure 5H**, P = 0.06, Mann-Whitney test). Together, these data indicate that the observed reduction in luminal γT3 in AVN-treated animals is driven by a microbiota modulation of the host γT3 absorption. These insights from *in vitro* and *in vivo* experiments close a mechanistic link between antibiotic-induced microbiome perturbation, altered γT3 bioavailability, and IDO1 downregulation.

### Validating Lachnospiraceae and Oscillospiraceae link to IDO1 in IBD patients

Finally, we tested whether the associations between Lachnospiraceae/Oscillospiraceae and colonic IDO1 expression observed in mice could also be detected in humans. We first analyzed our allo-HCT patient cohort, in which stool KYN/TRP ratios served as a proxy for gut IDO1 activity. LASSO regression on original and scaled abundances at the ASV, genus, and family levels did not identify any microbial predictors of stool KYN/TRP (data not shown). Similarly, direct correlation analysis between Lachnospiraceae abundance and stool KYN/TRP was not significant (Pearson correlation, R² = 0.0207, P = 0.84). These negative findings were likely influenced by the relatively small sample size (n = 92), the high dimensionality of the microbiome data, and the complex allo-HCT setting, in which patients experience simultaneous fluctuations in immune and microbiota markers.

To overcome these issues, we turned to a publicly available dataset from patients with IBD. This group of disorders includes Crohn’s disease (CD) and ulcerative colitis (UC), characterized by excessive intestinal immune activation. The dataset included paired colonic biopsy RNA-seq data and 16S rDNA microbiome profiles (**Figure 6A**)^39^. We saw that IDO1 expression was significantly higher in biopsies from IBD patients compared with healthy controls (**Figure 6B**, P < 0.0001, Mann-Whitney test), consistent with previous reports of KP activation in IBD^40,41^. LASSO regression, incorporating both microbial and clinical variables, identified active inflammation at the biopsy site as the strongest predictor of IDO1 expression (**Figure 6C**). Importantly, after controlling for gut inflammation, we still identify several bacterial taxa associated with IDO expression: the top microbial predictor was an ASV classified as *Blautia* (Lachnospiraceae), which was positively correlated with IDO1. Additional positive predictors included *Flavonifractor* (Oscillospiraceae) and *Agathobacter* (Lachnospiraceae), confirming that members of both families are associated with higher IDO1 expression in the human gut mucosa.

**Figure 6.**
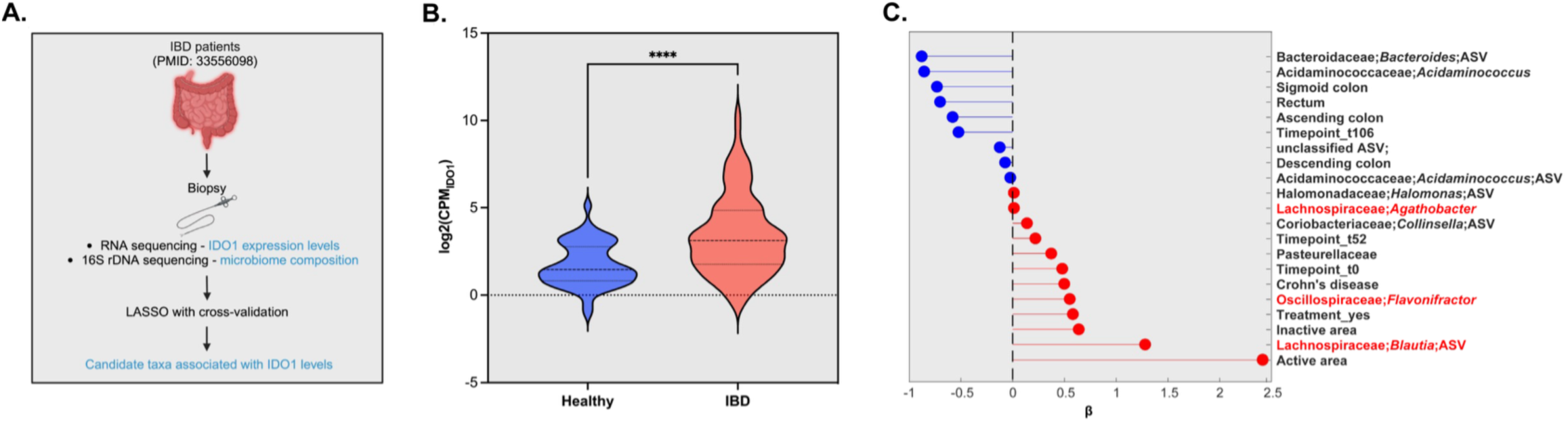
IDO1 expression levels in biopsies from a human inflammatory bowel disease (IBD) cohort correlate with the abundances of Lachnospiraceae and Oscillospiraceae bacteria. A) Schematic representation of the publicly available IBD patient cohort re-analyzed in this study. B) IDO1 levels are significantly higher in biopsies from IBD patients compared to those from healthy individuals. CPM: counts per million. P < 0.0001, two-tailed Mann-Whitney test. C) Lachnospiraceae and Oscillospiraceae correlate with IDO1 expression in biopsies from the IBD cohort. LASSO regression selected predictors of the IDO1 levels. Red circles represent taxa positively correlated with IDO1, while blue circles represent taxa negatively correlated. Taxa highlighted in red belong to the Lachnospiraceae and Oscillospiraceae families. β: LASSO coefficient. ASV: amplicon sequence variant. In violin plot, lines show median and quartile. See also **Figure S6**.

To validate these findings, we re-analyzed a second independent dataset^42^ comprising mucosal biopsies from patients with CD, UC, acute non-IBD related inflammation, and healthy controls (**Figure S6**). While IDO1 levels did not differ significantly between IBD patients (CD and UC) and healthy controls (**Figure S6A**, P > 0.99, Mann-Whitney test), stratification by inflammation status revealed significantly higher IDO1 expression in inflamed versus non-inflamed sites (**Figure S6B**, P < 0.05, Mann-Whitney test). In this dataset, the strongest microbial predictor of IDO1 was an OTU from the Lachnospiraceae family (**Figure S6C**), again showing a positive correlation with IDO1 expression.

These complementary analyses support the finding that Lachnospiraceae and Oscillospiraceae drive intestinal IDO1 expression in human cohorts independently from inflammation.

## DISCUSSION

The KP is a key regulator of host immunity, yet the mechanisms controlling its activity locally in the intestine remain poorly defined. Here, we demonstrate that intestinal KP activity can be downregulated by perturbing the microbiota with antibiotics. This robust finding led us to the discovery that the microbiota-mediated modulation of IDO1 expression works indirectly through the regulation of the absorption of the micronutrient γT3, independently of systemic inflammation.

At the molecular level, we concluded that γT3-mediated suppression of IDO1 depended on inhibition of HMG-CoA reductase and downstream mevalonate signaling (**Figure 5F**). This is in line with prior reports linking γT3 to regulation of the mevalonate pathway^36,43^. Mevalonate is a known inducer of NF-κB^44^, a transcription factor that promotes IDO1 transcription^45^. Pharmacological modulation of γT3 transport further demonstrated that host uptake is a critical determinant of its effects on IDO1 (**Figure 5F**). Together, these findings support a model in which commensal bacteria modulate the intestinal availability of a dietary γT3, thereby sustaining IDO1 expression in the gut epithelium.

Previous studies have implicated probiotic bacteria in modulating IDO1 activity^27–29^, yet the contribution of resident commensals and dietary metabolites has remained largely unexplored. Our data suggest that, under homeostatic conditions, Lachnospiraceae and Oscillospiracaeae from the commensal microbiota actively sustain intestinal IDO1 expression, counterbalancing the suppressive effects of dietary γT3. This mechanism is particularly relevant given that excessive KP activation has been linked to immune suppression in cancer^46^ and chronic inflammation^40^. Our findings highlight the importance of maintaining appropriately balanced IDO1 activity for intestinal immune homeostasis, and underscoring the nuanced role of KP regulation in health and disease.

More broadly, this work illustrates how microbiota-dependent control of nutrient absorption can shape local host metabolism, a microbiota-metabolism link with an established role in immune regulation, and whose dysregulation could lead to inflammatory diseases. Although the specific components of the microbiota secretome responsible for modulating γT3 absorption remain to be identified, our results open new avenues for defining postbiotic mediators and exploring dietary or microbiota-based strategies to fine-tune IDO1 activity. Such approaches may hold therapeutic potential in IBD, cancer, and other immune-mediated disorders where KP dysregulation contributes to pathology.

## RESOURCE AVAILABILITY

### Lead contact

Requests for further information and resources should be directed to and will be fulfilled by the lead contact, Joao B. Xavier.

### Materials availability

This study did not generate new unique reagents.

### Data availability

- Sequencing files generated by 16S rDNA sequencing have been submitted to NCBI and are publicly available (PRJNA1435869).
- Sequencing files generated by RNA sequencing of mouse tissues have been submitted to NCBI and are publicly available (PRJNA1435869).
- 16S rDNA abundance tables, metabolomic profiling data, IDO1 expression tables and other materials are available on Zenodo. (https://doi.org/10.5281/zenodo.18937393).
- Code used for analysis is available on GitHub (https://github.com/anadj-micro/Gut-microbiome-kynurenine-pathway-regulation.git).
- Any additional information required to reanalyze the data reported in this paper is available from the lead contact upon request.

## ACKNOWLEDGEMENTS

The authors would like to thank Dr. Eric G. Pamer and Dr. Carles Ubeda for providing us with the Lachnospiraceae and Oscillospiraceae strains, and Dr. Karuna Ganesh for providing us with MC-38 TGL cell line. We are also very grateful to Dr. Simon Grassmann for the help with germ-free mice. We would like to thank Molecular Microbiology Facility at Memorial Sloan Kettering Cancer Center (MSKCC) for DNA extraction and 16S rDNA processing, Molecular Cytology Core for FISH staining and slides processing, Proteomics and Metabolomics Core, RRID:SCR_027811, for metabolic profiling of patients’ stool and blood samples, and mouse blood. This work was supported by RYC2023-042654-I grant funded by MICIU/AEI/10.13039/501100011033 and by the ESF+, and by PID2024-160934OA-I00 grant funded by MICIU/AEI/10.13039/501100011033/FEDER/EU to A.Djukovic; the USA/NIH grants R00AI175599 to C. Liao, R01 CA266068, R01 AI196346 and P01 AI179406 to J. B. Xavier; Institutional Core Grant P30 CA008748 to Molecular Cytology Core at MSKCC; and Starr Cancer Consortium (grant SCC Award #: I13-0013).

## AUTHOR CONTRIBUTIONS

A.D. and J.B.X. conceived the study. A.D. performed mouse experiments, 16S rDNA qPCR, RT-qPCR, processing of the 16S rDNA amplicon data, statistical analysis of microbiome profiles and metabolomics data and *in vitro* experiments involving Lachnospiraceae and Oscillospiraceae strains. J.B.X. performed analysis of the allo-HCT patient cohort. C.Liao processed the publicly available datasets from IBD patients and performed statistical analysis. A.B.S. and C.L. performed MC-38 cell line *in vitro* experiments and helped with mouse experiments. H.Liu adapted analytical method for measuring tryptophan metabolites and plasma and stool of patients. H. Lees developed a pipeline for statistical analysis of metabolic profile in blood and stool of patients. R.J.F.R. designed the experimental plan, conducted the allo-HCT sample preparation, and was part of the data mining and analysis. J.C. supervised metabolic profiling of samples from allo-HCT patients and mouse plasma. A.D. and J.B.X. produced Figure 1. A.D. produced Figures 2-6. All co-authors read and contributed to the manuscript.

## DECLARATION OF GENERATIVE AI AND AI-ASSISTED TECHNOLOGIES

During the preparation of this work, the authors used ChatGPT in order to correct grammar and improve readability of the text. After using this tool, the authors reviewed and edited the content as needed and take full responsibility for the content of the publication.

## SUPPLEMENTAL INFORMATION

**Document S1**. Figures S1-S6.

**Table S1. Results of the microbiome composition comparison between responders and non-responders.** Related to **Figure 4**.

## STAR METHODS

### EXPERIMENTAL MODEL AND STUDY PARTICIPANT DETAILS

#### Human participants

Data obtained from human participants analyzed in this study have been previously published and are publicly available. The accession numbers for the datasets are listed in the key resources table. Metabolic profiling of stool and plasma of allo-HCT patients generated in this study is publicly available and accession number is listed in key resources table. This study re-analyzed two previously published datasets from IBD patients. The accession numbers for the datasets are listed in the key resources table.

#### Mouse model

Mice used in this study were C57BL/6J specific pathogen-free mice purchased from The Jackson Laboratories. Animals were 6-8 weeks old females. During the study, mice were mostly single-housed (except for the germ-free mouse experiments) in autoclaved cages with *ad libitum* access to autoclaved and acidified reverse osmosis water (pH, 2.5 to 2.8) and irradiated feed (LabDiet 5053, PMI, St Louis, MO). The animal holding room was maintained at 72 ± 2 F (21.5 ± 1 C), relative humidity between 30% and 70%, and a 12:12 hour light:dark photoperiod. For confirming the role of the microbiome in regulation of the KP we used germ-free C57BL/6J mice. In these experiments we used both males and females. Mice were co-housed in groups in autoclaved isolator cages with *ad libitum* access to autoclaved water and feed. Animal use is approved by Memorial Sloan Kettering Cancer Center’s IACUC (protocol # 18-03-003). The institution’s animal care and use program is AAALAC-accredited and operates in accordance with the recommendations provided in the *Guide*.

#### Bacterial strains

Bacterial strains used in this study were gift from Dr. Eric G. Pamer (*C. eutactus*, *B. massiliensis*, *R. intestinalis*), Dr. Carles Ubeda (*Flavonifractor spp*.), and purchased from DSMZ (*I. butyriciproducens* DSMZ 26588). All strains were routinely grown on Columbia Blood Agar (CBA) plates inside of anaerobic chamber at 37°C for 48 hours or more, as needed. Anaerobic conditions inside the anaerobic chamber were maintained with a gas mix containing 5% carbon dioxide, 7.5% hydrogen and nitrogen as a balance. Hydrogen levels were maintained above 3%.

#### Cell line

Cell line used in this study was MC-38 TGL colorectal cancer cell line gift from Dr. Karuna Ganesh. Cells were routinely grown in high-glucose DMEM with 10% fetal bovine serum (FBS) and 1% penicillin G-streptomycin at 37°C and 5% CO_2_.

## METHOD DETAILS

### Mouse experiments

#### Immune vs. microbiome-driven KP regulation

Mice were divided in three groups: untreated control, antibiotic treated group and LPS treated group. Each group had three animals. After initial period of acclimatation, animals were randomly assigned to groups and single-housed. Antibiotic-treated animals received a cocktail of ampicillin (0.5 g/L), vancomycin (0.5 g/L) and neomycin (1 g/L) for one week in drinking water. The water was changed once during treatment to preserve the activity of antibiotics. Untreated and LPS-treated animals received regular water. LPS-treated animals received LPS (15μg/animal in 0.9% NaCl) intraperitoneally. The day after LPS injection and two days after antibiotic cessation, feces were collected from all the groups for 16S rDNA profiling, kept on ice until reaching the lab, and then stored at -80°C until further processing at the Molecular Microbiology Facility at MSKCC. After feces collection, animals were fasted for 6 hours, and blood was collected for tryptophan metabolic profiling by performing a mandibular vein punction with 4mm lancet. Lancets were changed between animals. Blood was collected into K2 EDTA coated tubes that were inverted several times and kept on ice until reaching the lab. There, tubes were centrifuged at 3000g for 15 minutes at 4°C. Supernatant was collected with Pasteur pipette and transferred to a new 1.5ml tube. Samples were frozen and kept at - 80°C until further processing at the Cell Metabolism Core. After blood collection, all the animals were euthanized and approximately 30mg of the small intestine was collected in 500μl of RNA later. After 24 hours at 4°C, the RNA later was removed and tissue was stored at -80°C until further processing for RNA extraction and sequencing.

#### Microbiome recovery experiments

For studying the effect of microbiome recovery on the KP, mice were treated with the cocktail of ampicillin, vancomycin and neomycin as described previously and then subjected to different microbiome-recovery strategies. One group, the fecal microbiome transplant (FMT) group, was treated with AVN and then orally gavaged with fecal slurry from untreated animals. Briefly, the day after antibiotic cessation fecal pellets from untreated animals (3-5) were resuspended in 1.5ml of pre-reduced phosphate buffer saline (PBS), and 100μl of this fecal slurry was administered via oral gavage to each AVN-treated animal. This was repeated for two more days. Fecal pellets for 16S rDNA profiling and proximal colons for IDO1 profiling were collected the day after the last gavage as described previously. The second group (spontaneous recovery group) was treated with AVN and then left to recover for two weeks. After this recovery period, fecal pellets and colons were collected as described previously for further analysis. The AVN-treated and untreated groups served as controls.

#### Germ-free mice experiments

Mice were co-housed in isolator cages for the period of the experiment. Two independent experiments were performed. One was performed on male and another on female animals. Mice were randomly assigned to untreated or AVN-treated group. AVN group received AVN cocktail in drinking water as described previously. The day after antibiotic cessation animals were euthanized and colon samples were harvested, as described previously, for IDO1 profiling. In order to confirm the sterility, fecal pellets were collected before and after the experiments, resuspended in pre-reduced PBS and plated on CBA plates inside anaerobic chamber.

#### Microbiome library generation

To generate a library of microbiome compositions, mice were treated with different antibiotics in the drinking water. The treatment lasted for one week, and water with antibiotics was changed once during the treatment to preserve the activity of the antibiotics. The used antibiotics and their concentrations were: ampicillin (0.5g/L), vancomycin (0.5g/L), a cocktail of ampicillin, vancomycin and neomycin as described previously, metronidazole (0.25g/L) and ciprofloxacin (0.2g/L). Untreated animals served as control. The day after the antibiotic cessation fecal pellets were collected for 16S rDNA profiling. Animals were euthanized and colon tissue samples were collected, as described previously, for the IDO1 expression analysis.

#### Administration of probiotic strains

To investigate the role of Lachnospiraceae and Oscillospiraceae on IDO1 expression, mice were treated for a week with the AVN cocktail, as described previously, followed by the gavaging with the probiotic mix. AVN-treated and untreated animals served as control. All animals received normal, non-acidified water for drinking once the antibiotic treatment was removed, to allow for better survival of the probiotic mix and successful gut colonization.

For three consecutive days starting the day after antibiotic cessation, animals were orally gavaged with probiotic mix containing five representative Lachnospiraceae and Oscillospiraceae strains. Animals were gavaged as described previously^47^. Briefly, animals received 200μl of NaHCO_3_ to neutralize the acidity of the stomach. This was followed by 100μl of probiotic mix in PBS 20% glycerol, or just PBS 20% glycerol for the AVN group. The third day of gavaging mice were fasted for 4 hours before gavage to maximize the chances for successful gut colonization by probiotic strains.

Probiotic mix was prepared ahead and contained *R. intestinalis, B. massiliensis, C. eutactus*, *Flavonifractor spp.* and *I. butyriciproducens*. All five strains were plated on CBA plates inside the anaerobic chamber. Once grown, cultures were collected with loops, resuspended in PBS 20% glycerol and absorbance was measured. Next, all cultures were adjusted with PBS 20% glycerol until reaching the same absorbance. Equal volume of each culture was used for making a mix, which was then aliquoted in aliquots for each day of gavage and stored at -80°C.

The day after the last gavage, fecal pellets were collected for 16S rDNA profiling. Animals were euthanized and colon tissue was collected, as described previously, for analyzing IDO1 expression levels. Another piece of colon and cecal content were collected for metabolomic profiling. These samples were flash frozen in dry ice-ethanol bath and stored at -80°C until metabolites extraction.

### RNA extraction from mouse tissues

RNA extraction was performed using RNeasy Plus Mini Kit from Qiagen. Protocol was followed with modifications: a) pre-filled 2ml tubes with 1.5mm Zirconium beads were used for homogenization; b) for maximizing the yield 30μl of β-mercaptoethanol was added per 1ml of RLT buffer on the day of extraction. The quality of the extracted RNA was confirmed by gel electrophoresis and Nanodrop, and quantified using Qubit RNA HS Assay Kit. The obtained RNA was used for RNA sequencing commercially at GENEWIZ® NGS Services from Azenta Life Sciences (South Plainfield, NJ, USA), or for the analysis of the IDO1 expression levels via RT-qPCR.

### Intestinal RNA sequencing

RNA samples were quantified using Qubit 2.0 Fluorometer (Life Technologies, Carlsbad, CA, USA) and RNA integrity was checked using Agilent TapeStation 4200 (Agilent Technologies, Palo Alto, CA, USA). RNA sequencing libraries were prepared using the NEBNext Ultra RNA Library Prep Kit for Illumina using manufacturer’s instructions (NEB, Ipswich, MA, USA). Briefly, mRNAs were initially enriched with Oligod(T) beads. Enriched mRNAs were fragmented for 15 minutes at 94°C. First strand and second strand cDNA were subsequently synthesized. cDNA fragments were end repaired and adenylated at 3’ends, and universal adapters were ligated to cDNA fragments, followed by index addition and library enrichment by PCR with limited cycles. The sequencing library was validated on the Agilent TapeStation (Agilent Technologies, Palo Alto, CA, USA), and quantified by using Qubit 2.0 Fluorometer (Invitrogen, Carlsbad, CA) as well as by quantitative PCR (KAPA Biosystems, Wilmington, MA, USA).

The sequencing libraries were multiplexed and clustered onto a flowcell on the Illumina HiSeq instrument according to manufacturer’s instructions. The samples were sequenced using a 2×150bp Paired End (PE) configuration. Image analysis and base calling were conducted by the HiSeq Control Software (HCS). Raw sequence data (.bcl files) generated from Illumina HiSeq was converted into fastq files and de-multiplexed using Illumina bcl2fastq 2.20 software. One missmatch was allowed for index sequence identification.

After investigating the quality of the raw data, sequence reads were trimmed to remove possible adapter sequences and nucleotides with poor quality using Trimmomatic v.0.36. The trimmed reads were mapped to the reference genome available on ENSEMBL using the STAR aligner v.2.5.2b. The STAR aligner uses a splice aligner that detects splice junctions and incorporates them to help align the entire read sequences. BAM files were generated as a result of this step. Unique gene hit counts were calculated by using feature Counts from the Subread package v.1.5.2. Only unique reads that fell within exon regions were counted. After extraction of gene hit counts, the gene hit counts table was used for downstream differential expression analysis. Comparison of gene expression between the groups of samples was performed using DESeq2.

### RT-qPCR for IDO1 expression

IDO1 expression in intestinal tissue and cell culture was determined using ΔΔCt method. Briefly, 2- 2.5μg of RNA extracted from colonic tissue/MC-38 cells was subjected to cDNA generation using SuperScript IV VILO kit. Samples were diluted in duplicates to reach equal concentration. Next, all the samples were treated with DNase as per manufacturer’s protocol, after which one set of duplicates was incubated with reverse transcriptase mix, while another set was treated with mix lacking reverse transcriptase. These samples were later used as no reverse transcriptase controls (NRT) in RT-qPCR. The obtained cDNA was diluted 1:1 in miliQ water to ensure enough volume for the RT-qPCR reactions, and then used to prepare TaqMan RT-qPCR reactions. For each reaction 10μL of the TaqMan Master Mix was combined with 1μL of TaqMan probe, 7μL of miliQ water and 2μL of cDNA. We used commercial IDO1 (Mm00492590_m1) and HMBS (Mm01143545_m1) probes, with HMBS serving as endogenous control. HMBS gene was selected after initial tests in which the Ct values of HMBS, ACTB, RPLP2 and GAPDH genes were compared between untreated and AVN-treated animals, reveling that HMBS gene showed no change in Ct values between tested groups. Each sample was run in triplicate. For NRT controls, samples subjected to cDNA generation with the SuperScript mix lacking reverse transcriptase were used as templates. For negative controls, water was used instead of cDNA. Cycling conditions were 50°C for 2 minutes, 95°C for 10 minutes and 45 cycles of 95°C for 15 seconds, followed by 60°C for 1 minute. IDO1 expression of each sample was normalized by its own HMBS value and mean value of the control samples.

### FISH staining and slides processing

#### Tissue fixation

For localizing IDO1 expression, colons were cut into 2 pieces. Fecal content was removed by gently pressing the colon with forceps. Pieces were laid into histology cassette next to each other, and returned to the round shape with the help of the forceps. Cassettes were put into a jar with freshly made 4% paraformaldehyde (PFA) and left over night at 4°C. Next day, cassettes were washed twice in water, submerged into 70% ethanol, and taken to Molecular Cytology Core for paraffin embedding and slicing. There tissue samples were processed following Leica HistoCore Pegasus protocol, and embedded with Leica HistoCore Pegasus Parablocks. Paraffin-embedded tissue was sectioned with Leica RM2255 at 5μm and kept at 4°C.

#### Staining

Samples were loaded into Leica Bond RX, baked for 30 minutes at 60°C, dewaxed with Bond Dewax Solution (Leica, AR9222), and pretreated with EDTA-based epitope retrieval ER2 solution (Leica, AR9640) for 15 minutes at 95°C. The probe (Advanced Cell Diagnostics, ready to use, no dilution) was hybridized for 2 hours at 42°C. Mouse PPIB (ACD, Cat# 313918) and dapB (ACD, Cat# 312038) probes were used as positive and negative controls, respectively. The hybridized probes were detected using RNAscope 2.5 LS Reagent Kit – Brown (ACD, Cat# 322100) according to manufacturer’s instructions with some modifications (DAB application was omitted and replaced with Fluorescent CF594/Tyramide (Biotium,92174) for 20 minutes at room temperature (RT)). After the run was finished, slides were washed in PBS and incubated with 5μg/mL 4’,6-diamidino-2-phenylindole (DAPI) (Sigma Aldrich) in PBS for 5 minutes, rinsed in PBS and mounted in Mowiol 4–88 (Calbiochem). Slides were kept overnight at -20°C before imaging.

After the slides were scanned, the coverslips were removed and slides were loaded into Leica Bond RX for double immunofluorescence (IF) staining. Samples were pretreated with EDTA-based epitope retrieval ER2 solution (Leica, AR9640) for 20 minutes at 100°C. The 4-plex IF staining and detection were conducted sequentially. The primary antibodies against MHCII/A488 (2.5μg/mL, BD Pharmingen, 556999), CD3/C430 (0.01μg, Abcam, ab135372), E-cadherin (0.0194μg, BD Biosciences,610181) or CD45/A647 (0.039μg/mL, Abcam, ab10558) were incubated for 1 hour at RT. For rabbit antibodies, Leica Bond Polymer anti-rabbit HRP (included in Polymer Refine Detection Kit (Leica, DS9800) was used, for the rat antibody and the mouse antibody, a rabbit anti-rat (Vector Lab, VA-4000) or a rabbit anti-mouse (Abcam, ab133469) secondary antibody were used as linkers for 8 minutes, before the application of the Leica Bond Polymer anti-rabbit HRP for 8 minutes at RT. After that, Alexa Fluor tyramide signal amplification reagents (Life Technologies, B40953, B40958) or CF® dye tyramide conjugates (Biotium, 92172, 96053) were used for IF detection. After the run was finished, slides were washed in PBS and mounted in Mowiol 4–88 (Calbiochem). Slides were kept overnight at -20°C before imaging.

#### Scanning and processing

Slides were scanned with Pannoramic P250 Flash Scanner (3DHistech, Hungary) using a 40x/0.95NA objective lense. Two sections per animal were drawn and exported as .tif files using Slide Viewer (3DHistech, Hungary). Images were then analyzed using ImageJ/FIJI (NIH, USA). DAPI was thresholded and masked in order to count cells. IDO1 foci counts per cell were then obtained using maxima finding on the images. Thresholding and measurement of fluorescence intensity was used to quantify the E-cadherin and IDO1. For all the quantifications, average values between two sections were calculated to obtain a single measurement per animal.

### Fecal DNA extraction and amplicon sequencing

DNA extraction and library preparation of the fecal samples was done at the Molecular Microbiology Facility at MSKCC. Briefly, fecal DNA was extracted and the 16S rDNA gene was amplified using a previously described protocol^48^. The QIAseq 1-step Amplicon Library kit (180419) was used for generating libraries that were later quantified, normalized and sequenced using MiSeq Reagent kit V3 (MS-102-3001).

### 16S rDNA qPCR

For assessing the total bacterial load in fecal samples, qPCR against standard curve was used to determine 16S rDNA copy number. For this purpose, the PowerUP qPCR kit (4367659) was used. Briefly, for each sample, 20μL PCR triplicates were prepared with each containing 2μL of the DNA used as template, 10μL of mix provided by the manufacturer, and 1μL of forward and reverse primers at the final concentration of 0.5μM. We used the primer pair 27F/338R to amplify the V1–V2 region of the 16S rRNA gene (F- AGAGTTTGATCMTGGCTCAG; R- TGCTGCCTCCCGTAGGAGT). To complete the volume of the reaction, 6μL of water was added. A PCR product of the 16S rRNA gene from *Enterococcus faecium* ATCC 700221 strain was used for obtaining a standard curve by amplifying its 16S rRNA gene and purifying the product. The copy number of the PCR product was determined on the basis of its concentration and 16S rDNA sequence. A standard curve was obtained by using 10-fold dilutions.

Cycling conditions of the qPCR were 50°C for 2 minutes, 95°C for 2 minutes and 40 cycles of 95°C for 15 seconds, 56°C for 15 seconds and 72°C for 60 seconds. By extrapolating results by looking at the ones obtained from standard curve samples, the number of 16S rRNA genes was determined for each sample. The final number of 16S rRNA genes per 1 gram of fecal sample was calculated by multiplying the number of 16S rDNA molecules obtained by qPCR with the DNA elution volume after DNA extraction and dividing this number by the weight of the fecal pellet from which DNA extraction was performed.

### Processing of the 16S rDNA amplicon data

Sequence processing was done following the Divisive Amplicon Denoising Algorithm (DADA2) tutorial with an in-house script. Briefly, after demultiplexing, reads were trimmed to the first 180 base pairs or the first point with a quality score Q < 2 and removed if they contained ambiguous nucleotides (N) or if two or more errors were expected on the basis of the quality of the trimmed reads. Paired reads were merged and chimaeras were removed. ASVs were identified using DADA2 and classified against the SILVA v.138 database^49^.

### Diversity calculation

In order to evaluate the richness and evenness of gut microbiome in the fecal samples from mice treated with different antibiotic treatments and those subjected to microbiome recovery regimes, we determined α-diversity by calculating Inverse Simson Index using formula i=1/∑Nj=1 *p*^2^*ij*, where N is total number of ASVs in the particular sample i and p are the relative abundance of the j-th ASV out of N.

### IBD patients’ data processing

For the patient cohort in PMID 33556098, 16S amplicon sequencing data was processed using the same pipeline described previously^31^. Sample metadata and gene expression data were downloaded from the original study. For the patient cohort in PMID 27694142, the 16S OUT count table, taxonomic classifications, gene expression data, and clinical metadata were downloaded from the original study. No additional processing was required. For both datasets, samples with fewer than 1,000 sequences were excluded from the analysis.

### Metabolites extraction

#### Cecal content/feces

Approximately 18mg (18-21mg) of cecal content or feces (16-22mg) was subjected to metabolites extraction. All the cecal/feces samples were extracted on the same day with the extraction solvent prepared on the day of extraction, in hand-washed glassware that was pre-rinsed with the solvent. The extraction solvent was 2:1 acetone:isopropanol (LCMS grade). 300μL of the solvent was added to each sample, and the samples were vortexed twice for 15 seconds intervals. The supernatant was passed to another tube, and the extraction step was repeated once again. All the supernatant was then centrifuged at 13 500rcf for 10 minutes at 4°C. 300μL of the supernatant was transferred to tubes provided by General Metabolics and stored at -80°C until shipping.

#### Cecal tissue

Approximately 15mg (14-15mg) of cecal tissue was subjected to metabolites extraction. All the samples were extracted on the same day with the extraction solvent prepared on the day of extraction, in hand-washed glassware that was pre-rinsed with the solvent. The extraction solvent was 40:40:20 acetonitrile:methanol:water (LCMS grade). 300μL of the solvent was added to each sample, together with 4-5 metallic beads, and samples were homogenized at full speed for one minute at TissueLyser. Additional 1mL of the cold extraction solvent was added to each sample followed by a short vortex. Samples were stored at -20°C for 18 hours, after which they were centrifuged at 4 000rpm for 10 minutes at 4°C. 300μL of the supernatant was transferred to the tubes provided by General Metabolics and stored at -80°C until shipping.

#### Bacterial supernatants

For assessing γT3 production/consumption by Lachnospiraceae and Oscillospiraceae strains, strains were grown anaerobically, and supernatants were collected for metabolite extraction. For the media, we used filtered cecal content from untreated mice and spiked it with a small amount of γT3, for accuracy purposes. Briefly, cecal content was harvested from euthanized mice, put immediately to dry ice and stored at -80°C. On the day of the experiment, 10x weight of water was added to the tube, vortexed vigorously and centrifuged at maximum speed for 5 minutes. The supernatant was filter sterilized, first through 0.45μm filter, and then again through 0.2μm filter. γT3 diluted in ethanol and DMSO was added to this cecal media to reach the final concentration of 70nM, which was double the usual concentration of cecal content, as per our preliminary analysis. The media was pre-reduced inside the anaerobic chamber for 2 hours, after which a single colony of each strain was used to inoculate 300μL of the media. From here, 100μL was centrifuged at maximum speed for 5 minutes and filtered through 0.2μm filter. 30μL of this filtrate, representing timepoint 0, was stored at -80°C for metabolites extraction. The leftover 200μL was incubated for 48 hours inside the anaerobic chamber, after which an aliquot was taken and processed the same way as for the timepoint 0.

All the bacterial supernatant samples were extracted the same day with the extraction solvent prepared on the day of extraction, in hand-washed glassware that was pre-rinsed with the solvent. The extraction solvent was 80% methanol (LCMS grade). Samples were thawed, vortexed for 15 seconds, and centrifuged briefly. 20μL of the sample was transferred to a 96-well plate, and 180μL of the solvent was added to each sample. Plates were caped, vortexed vigorously for 15 seconds, and then incubated for 1 hour at 4°C. Next, the plate was centrifuged at 4 000rpm for 30 minutes at 4°C in a pre-cooled centrifuge. 100μL of the supernatant was transferred to a new plate, and from here 50μL of the supernatant was transferred into tubes provided by General Metabolics. Tubes were stored at - 80°C until shipping.

#### Allo-HCT stool, plasma and mouse plasma

Extraction of the metabolites from allo-HCT patient cohort, as well as from the mouse plasma samples for distinguishing immune- from microbiome-driven regulation of KP was performed at Proteomics and Metabolomics Core at MSKCC.

Human and mouse plasma samples were thawed on wet ice for 1 hour and vortexed for 10 seconds. A 100μL aliquot of each sample (50μL for mouse plasma) was transferred into pre-labeled 1.5mL microcentrifuge tubes containing 400μL of extraction solvent (100% methanol) supplemented with internal standards (ISTDs). Samples were vortexed for 5 seconds and incubated overnight at -80°C to facilitate protein precipitation. The following day, extracts were incubated on wet ice for 1 hour, vortexed briefly, and centrifuged at 20,000*g* for 20 minutes at 4°C. The resulting supernatant (450μL) was transferred into a new microcentrifuge tube and dried under vacuum using a Genevac EZ-2 Elite evaporator. Dried extracts were resuspended in 100μL of 0.2% formic acid in water, vortexed, and incubated on ice for 20 minutes. Samples were then clarified by centrifugation at 20,000*g* for 20 minutes at 4°C. Pooled quality controls (PQC) were prepared by combining 5μL aliquots from each sample.

Approximately 120mg (wet weight) of human fecal material was weighed into 2mL bead tubes containing 1.4mm ceramic beads (Omni International). Samples were extracted using 80:20 methanol:water supplemented with internal standards (ISTDs) to a final concentration of 100mg/mL. Homogenization was performed using a Bead Ruptor 24 (Omni International), and the resulting slurries were stored at -80°C overnight. Following storage, the extracts were thawed, vortexed, and centrifuged at 20,000*g* for 20 minutes at 4°C. An aliquot of 100μL of the supernatant was transferred to a 96-well collection plate (Waters) and dried under vacuum using a Genevac evaporator. Dried extracts were reconstituted with 100μL of 0.2% formic acid in water and vortexed at 400 rpm at 4°C for 20 minutes. Samples were then centrifuged at 1349*g* for 30 minutes at 8°C and transferred to an Agilent injection plate for analysis. Pooled quality controls (PQC) were prepared by combining 5μL aliquots from each sample.

### Metabolites profiling

#### Mouse feces, cecal content, tissue, and bacterial supernatants

Samples were shipped overnight on dry ice to General Metabolics. Metabolic extracts of cecal content, tissue, and bacterial supernatants samples were diluted by a factor of 1-to-50 in a chilled solution of 80% methanol in water (v/v) prior to analysis. A set of positive control samples (“pSS_spike” and “blank_spike”) were acquired along with the biological samples. These samples were spiked at a final concentration of 100nM of gT3. A pooled study sample (pSS) was prepared and injected periodically throughout the full course of the data acquisition run to facilitate assessment of technical reproducibility. For fecal extracts, samples were thawed and diluted 1:1000 in a chilled solution of 80% methanol in water (v/v) prior to analysis. As for the previous run, pooled study sample (pSS) was prepared and injected periodically throughout each batch.

Metabolome profiles of the sample extracts were acquired using flow-injection mass spectrometry. The method described here is adapted from Fuhrer et al. 2011^50^. The instrumentation consisted of an Agilent 6550 iFunnel LC-MS Q-TOF mass spectrometer in tandem with an MPS3 autosampler (Gerstel) and an Agilent 1260 Infinity II quaternary pump. The running buffer was 60% isopropanol in water (v/v) buffered with 1mM ammonium fluoride. Hexakis (1H, 1H, 3H-tetrafluoropropoxy)-phosphazene) (Agilent) and 3-amino-1-propanesulfonic acid (HOT) (Sigma Aldrich). The isocratic flow rate was set to 0.150mL/minute. The instrument was run in 4GHz High Resolution, negative ionization mode. Mass spectra between 50 and 1,000 m/z were collected in profile mode. 5μL of each sample were injected twice, consecutively, within 0.96 minutes to serve as technical replicates. The pooled study sample was injected periodically throughout the batch. Samples were acquired randomly within plates. Raw profile data were centroided, merged, and recalibrated using algorithms adapted from Fuhrer et al. 2011^50^. Putative annotations were generated based on compounds contained in the Human Metabolome Database, KEGG, and ChEBI databases using both accurate mass per charge (tolerance 0.001 m/z) and isotopic correlation patterns. LOESS normalization was applied on ion intensities table from cecal content, tissue and bacterial supernatant samples prior to statistical analysis to account for a modest temporal instrument drift in the pooled study sample.

#### Allo-HCT stool, plasma and mouse plasma

Samples were processed at Proteomics and Metabolomics Core at MSKCC. Samples were analyzed using reverse-phase liquid chromatography coupled to a TSQ Vantage triple quadrupole tandem mass spectrometer (Thermo Scientific) operating in positive electrospray ionization (ESI) mode. Chromatographic separation was performed on an Agilent 1260 Infinity binary pump system equipped with an ACQUITY UPLC HSS T3 column (2.1 × 100mm, 1.8μm, Waters), maintained at 35°C using a MayLab column oven. The mobile phases consisted of (A) 0.1% formic acid in water and (B) 0.1% formic acid in acetonitrile. The gradient elution program was as follows: 0-2 minutes, 0% B; 2-5 minutes, linear increase to 12% B; 5-7 minutes, linear increase to 70% B; 7-8.5 minutes, linear increase to 97% B; 8.5-11.5 minutes, hold at 97% B; followed by 3.5-minute re-equilibration time at initial conditions. The chromatographic parameters were: flow rate, 300mL/min; injection volume, 15µL; and column temperature, 35°C. Mass spectrometry settings were: capillary temperature, 300°C; vaporizer temperature, 350°C; sheath gas, 50; aux gas, 30; and spray voltage, 4000V. Data acquisition and analysis were performed using TraceFinder software (version 4.1, Thermo Scientific). Compound identities were confirmed by comparison with authentic standards.

### MC-38 model establishment

MC-38 TGL colorectal cancer cell line was gift from Dr. Karuna Ganesh. Cells were routinely grown in high-glucose DMEM with 10% FBS and 1% penicillin G-streptomycin. To establish an adequate model for measuring IDO1 expression, we grew the cells in standard 2-D conditions, and in low-attachment plates allowing for formation of 3-D cell spheroids. Briefly, 6-well plates were seeded with 3 x 10^5^ cells per well and left for 72 hours before RNA extraction.

For testing the inducibility of the IDO1 in MC-38 cell culture, cells were exposed to IFNγ. Briefly, 6-well plates were seeded with 4 x 10^5^ cells per well and left for a week to establish spheroids. After first 48 hours, media was exchanged and cells were left to grow for 4 more days. On the fourth day, media was exchanged again. Control cells received regular media, and another set of wells received media spiked with IFNγ (final concentration 1ng/mL). After 24 hours RNA was extracted.

### **γ**T3 effect on IDO1 levels in MC-38 cells

#### **γ**T3 alone

For testing the effect of γT3 on IDO1 expression levels in 3-D cultures of MC-38 cell line, cells were exposed to increasing concentration of γT3 (5μM, 25μM, 50μM, 75μM) for 24 hours. Media used was high-glucose DMEM with 10% dialyzed FBS (dFBS; for more control over the metabolic pathways) and 1% penicillin G-streptomycin. Briefly, 4 x 10^5^ cells were seeded per well in 6-well low attachment plates and left for a week to establish spheroids. After first 48 hours, media was exchanged and cells were left to grow for 4 more days. On the fourth day, media was exchanged again. Control cells received regular media with vehicle, and other wells received media spiked with γT3 in ethanol to reach previously mentioned final concentrations. The volume of the ethanol (vehicle) was the same between all the conditions. After 24 hours RNA was extracted.

#### **γ**T3 with probiotic secretomes

For testing the combined effect of γT3 and probiotic supernatants on IDO1 expression levels in 3-D cultures, cells were exposed to 25μM of γT3 in combination with secretomes from *R. intestinalis, B. massiliensis, C. eutactus*, *Flavonifractor spp.* and *I. butyriciproducens*. The media included dFBS as previously described. Briefly, 4 x 10^5^ cells were seeded per well in 6-well low attachment plates and left for a week to establish spheroids. After first 48 hours, media was exchanged and cells were left to grow for 3 or 4 more days. On the third or fourth day, media was exchanged again. Control cells received regular media with vehicle, γT3 group received γT3 at final concentration of 25μM, and other wells received γT3 (25μM) in combination with bacterial secretomes (see below). The volume of the ethanol was the same between all the conditions. After 24 or 48 hours RNA was extracted.

For preparation of bacterial secretomes, five Lachnospiraceae and Oscillospiraceae strains were grown on CBA plates inside the anaerobic chamber at 37°C for 48 hours. Cultures were collected with loops and used to inoculate 50mL of pre-reduced high-glucose DMEM with 10% dFBS and no antibiotics. After 48 hours of incubation, cultures were centrifuged, and supernatant was sterilized by passing it through 0.2μm filters. The obtained secretomes were mixed 1:1 with fresh high-glucose DMEM 10% dFBS 1% penicillin G-streptomycin and stored at -20°C. For each experiment, supernatants were thawed at 4°C over night, spiked with γT3, and added to MC-38 cells for periods of 24 or 48 hours, as described previously.

#### **γ**T3 with transport inhibitors and mevalonate

For testing the combined effect of γT3 and transport inhibitors or mevalonate on IDO1 expression levels in 3-D cultures, cells were exposed to 25μM of γT3 in combination with ezetimibe (an inhibitor of Niemann-Pick C1-Like 1 transporter), BLT-1 (a scavenger receptor BI inhibitor), or mevalonate (an HMG-CoA reductase product). 4 x 10^5^ cells were seeded per well in 6-well low attachment plates and left for a week to establish spheroids. After first 48 hours, media was exchanged and cells were left to grow for 4 more days. On the fourth day, media was exchanged again. Control cells received regular media with vehicle, γT3 group received γT3 at final concentration of 25μM, and other wells received γT3 (25μM) in combination with mentioned compounds. Final concentration for ezetimibe was 20μM, for the BLT-1 1μM, and for the mevalonate 2μM. These compounds were dissolved in DMSO, thus ethanol and DMSO were added to all the conditions reaching the equal volume. After 24 hours of incubation RNA was extracted.

### RNA extraction from cell culture

RNA extraction was performed using RNeasy Plus Mini Kit from Qiagen. For the 2-D cultures, media was aspirated and cells were washed with PBS, after which they were scraped and passed to a new tube, and resuspended in 350μL of the RLT buffer with β-mercaptoethanol. For the 3-D cultures, spheroids were collected into tube, centrifuged, washed with PBS, and resuspended in 350μL of the RLT buffer with β-mercaptoethanol before proceeding with the homogenization step. For homogenization, cells in RLT buffer were passed through QIAshredder column, after which the RNeasy Plus Mini Kit’s protocol was followed. The quality of the RNA was confirmed by Nanodrop and quantified using Qubit RNA HS Assay Kit. The extracted RNA was used for the RT-qPCR to analyze IDO1 expression levels, as described previously.

## QUANTIFICATION AND STATISTICAL ANALYSIS

### Analysis of allo-HCT patients’ data

We used the subset of stool samples selected from our previously published fecal microbiome atlas^30^ to span both biodiverse microbiota states and antibiotic-perturbed, low-diversity states dominated by a single taxon. Sample positions on the TaxUMAP embedding were obtained from the atlas coordinates, and each sample was colored according to the taxon of the most abundant amplicon sequence variant (ASV) in that stool sample. To assign this color, the highest-count ASV per sample was identified from the ASV count table and matched to its taxonomy annotation and corresponding color. Of the 96 samples with paired stool and plasma metabolite measurements, 4 did not have TaxUMAP coordinates (1107A, 1297D, FMT.0009O, and FMT.0079J) and were excluded from the embedding-based analyses, leaving 92 samples for the displayed cohort. Stool and plasma kynurenine-to-tryptophan (KYN/TRP) ratios were calculated for each sample by dividing measured kynurenine by measured tryptophan. Associations between neopterin and KYN/TRP were tested using Spearman rank correlation with pairwise complete observations. After TaxUMAP filtering, the plasma analysis included the 46 samples with available plasma neopterin measurements, whereas the stool analysis included all 92 remaining samples. For figure display, both correlation panels were shown on log-scaled axes; small offsets were added only to permit plotting of undetectable values on the logarithmic x-axis.

### RNA-seq analysis

For comparing the KP activity locally, in the gut, DESeq2 counts table was used to normalize each sample based on the geometric mean. Briefly, a pseudo-reference sample was established by calculating geometric mean for each of the rows. Next, the ratio of each sample to the reference sample is calculated, followed by the size factor calculations by computing the mean of each of the ratios. Next, counts were divided by their sample size factor. Negative binomial test was run to compute p-values for each gene when comparing samples from untreated and AVN-treated, and untreated and LPS-treated mice (“MATLAB” function *nbintest*). Next, the Benjamini-Hochberg adjusted p-values and log2 fold changes were calculated. A gene enrichment analysis was performed by using hypergeometric probability density function (“MATLAB” function *hygepdf*), followed by the multiple hypothesis correction (“MATLAB” function *mafdr*).

### LASSO regression

In order to determine which taxa and metabolites explain IDO1 expression levels in mice patients, we applied LASSO regression (“MATLAB” function *lasso*). For identification of the taxa, all samples with less than 1,000 sequences were removed from analysis. A wide table containing ASV, genus and family relative or absolute (relative multiplied by the total 16S rDNA in the sample) abundances was constructed, and taxa were used as predictors. The IDO1 expression in the corresponding colon sample was used as a response variable. The taxa abundances were min-max scaled to avoid giving more weight to the more abundant taxa. For the identification of metabolites of interest, all the identified metabolites were included in the analysis. As with taxa, min-max standardization was performed. To determine the best value of the regularization parameter, a five-fold cross-validation was performed during the model fitting. The relative weight of the lasso penalties was determined by setting the parameter alpha to 1. Taxa/metabolites selected by lasso as predictors of the IDO1 were used to fit a linear regression model (“MATLAB” function *fitlm*).

For IBD cohorts, ASV (or OTU), genus and family relative abundances were used as predictors. The IDO1 expression values were converted to log2-transformed counts per million and used as a response variable. The taxa abundances were min-max scaled to avoid giving more weight to the more abundant taxa. LassoCV function in Python was implemented, with 5-cross validation, and n_alphas = 1,000.

### Pearson correlation

For analyzing the correlation between microbiome diversity, represented by Inverse Simpson Index, or microbiome load, represented by total 16S rDNA counts, and IDO1 levels in colon, we used Pearson correlation (“MATLAB” function *corr*, specification *pairwise*). The same was used to correlate Lachnospiraceae abundances in allo-HCT cohort with the stool KYN/TRP ratio.

### Group comparisons

For statistical comparisons between two experimental groups, we used two-tailed Mann-Whitney test implemented in Prism, except for Figures 5A and 5B where previous studies already suggested expected signature, which is why we used one-tailed Mann-Whitney. For comparison between more than two experimental groups, we used regular one-way ANOVA, also implemented in Prism. Figure legends contain information on statistical tests and significance cutoffs for each of the panels.

