## Supplemental Figures for "Gut Commensals Regulate the Intestinal Kynurenine Pathway"

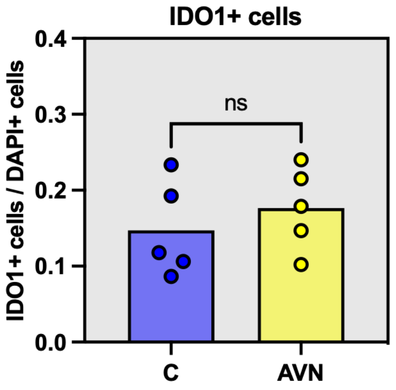


**Supplementary Figure 1. The number of IDO1-positive cells in colonic cross-sections remains unchanged after antibiotic cocktail treatment.** Fraction of IDO1-positive cells relative to the total number of cells, as measured by DAPI staining, in control (C) and antibiotic cocktail-treated animals (AVN). Ns: not significant, two-tailed Mann-Whitney test. Bars represent mean, and error bars represent the standard deviation. Related to **Figure 2**.

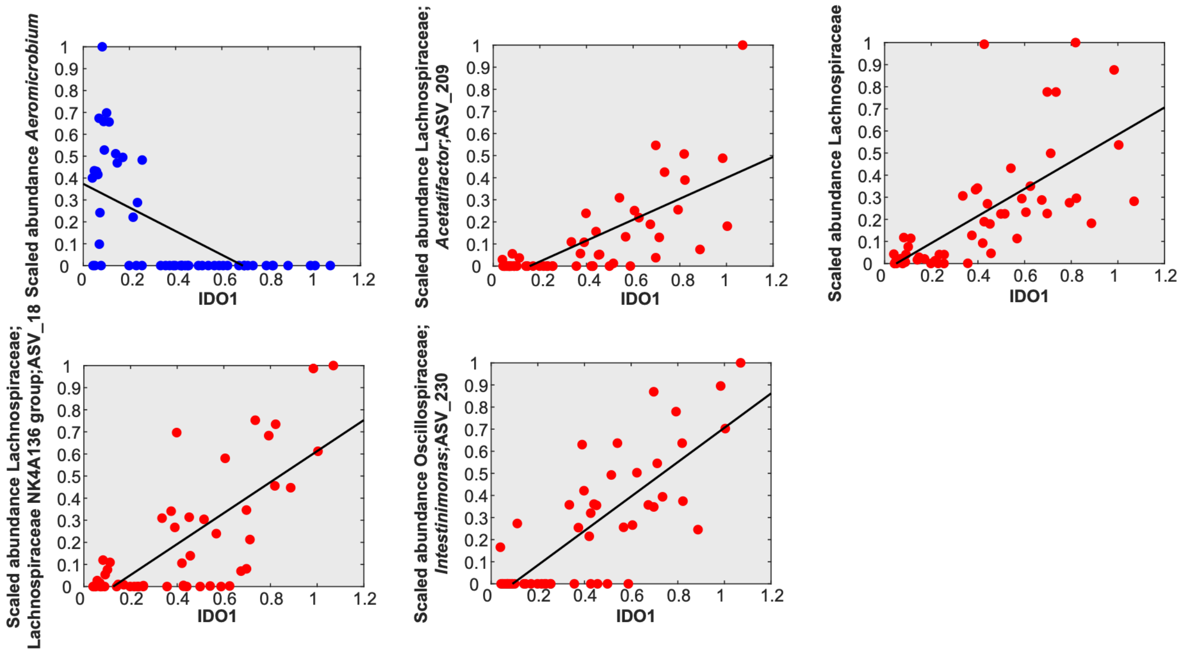


**Supplementary Figure 2. Correlation between scaled abundances of IDO1 predictors in fecal samples and the corresponding colonic IDO1 levels.** Each plot represents a single predictor. Predictors negatively correlated with IDO1 are shown in blue, while those positively correlated are shown in red. The regression line is plotted. Related to **Figure 3**.

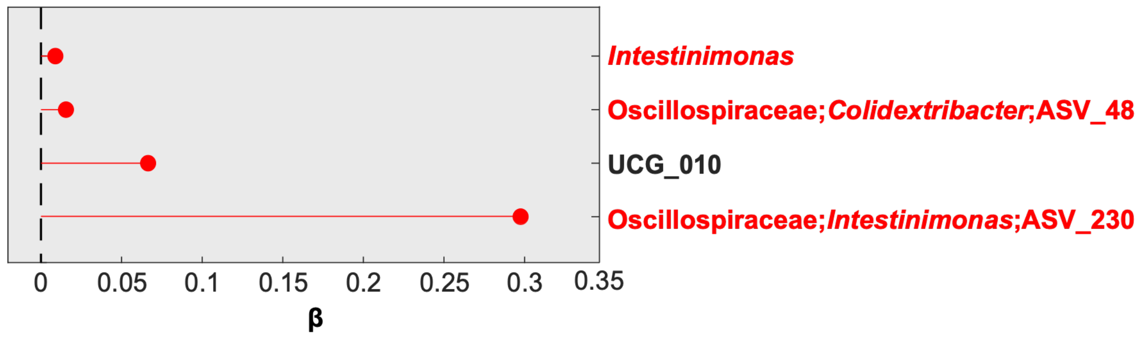


**Supplementary Figure 3. Taxa selected by LASSO regression as predictors of IDO1 levels in the colon when absolute abundances were used as input.** Red circles represent taxa positively correlated with IDO1. Taxa highlighted in red belong to the Lachnospiraceae and Oscillospiraceae families. β: LASSO coefficient. ASV: amplicon sequence variant. Related to **Figure 3**.

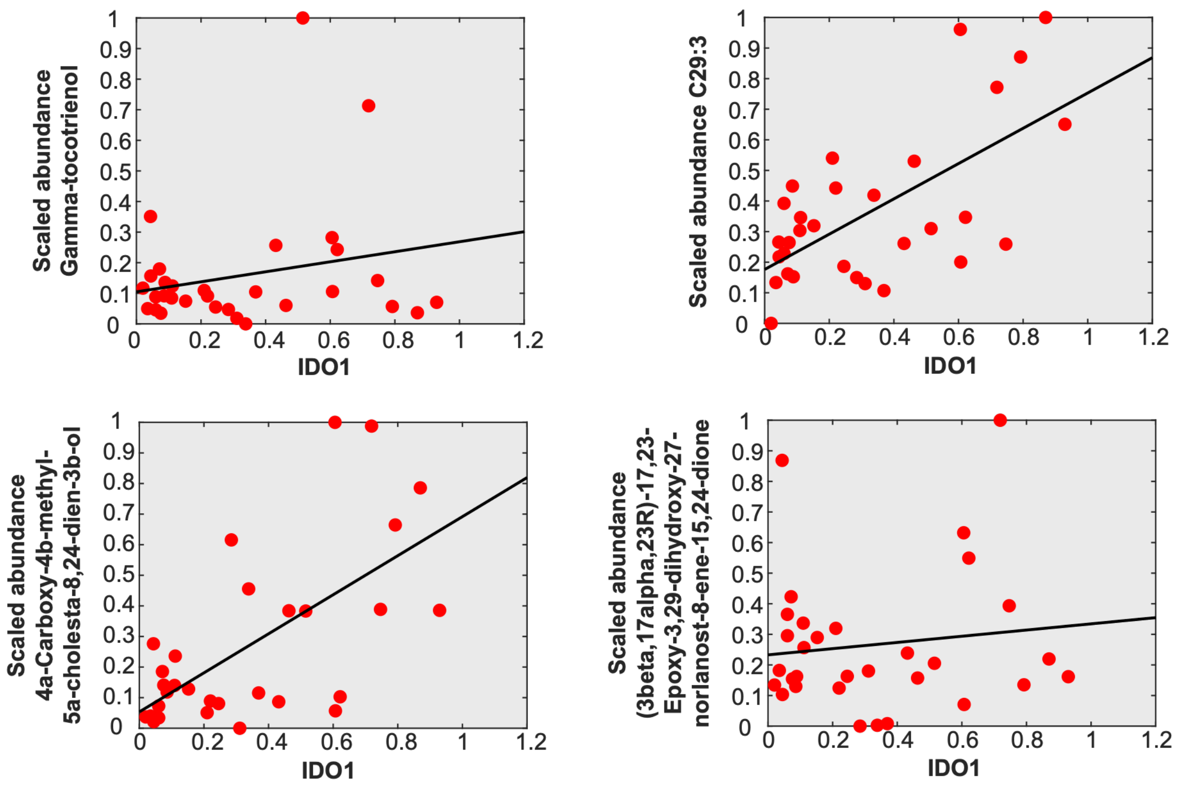


**Supplementary Figure 4. Correlation between scaled abundances of metabolites IDO1 predictors and the corresponding colonic IDO1 levels.** Each plot represents a single predictor. Predictors positively correlated with IDO1 are shown in red. The regression line is plotted. Related to **Figure 4**.

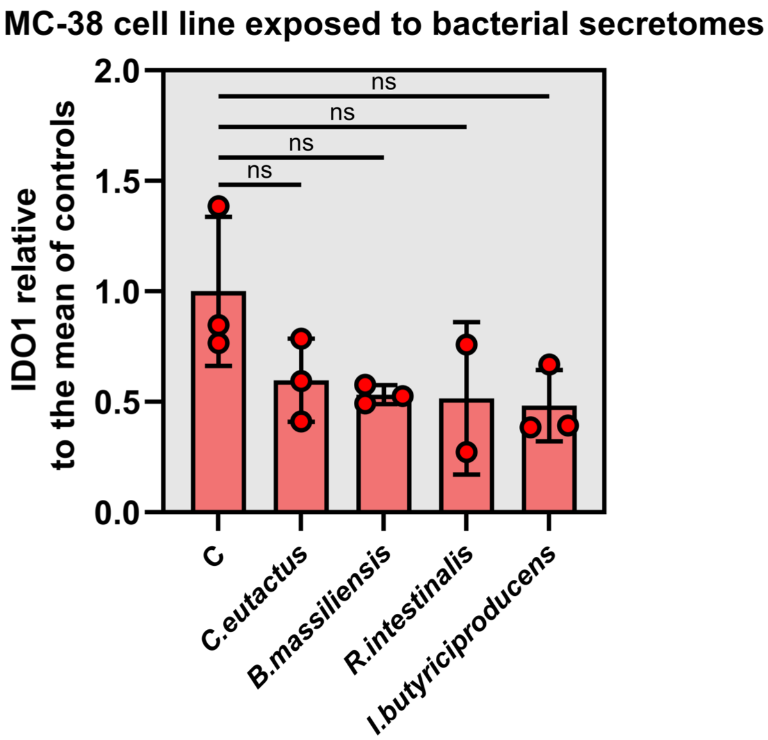


**Supplementary Figure 5. The IDO1 expression is unaffected by exposure to bacterial secretomes alone.** $P > 0.05$, one-way ANOVA. Bars represent mean, and error bars represent the standard deviation. Related to **Figure 5**.

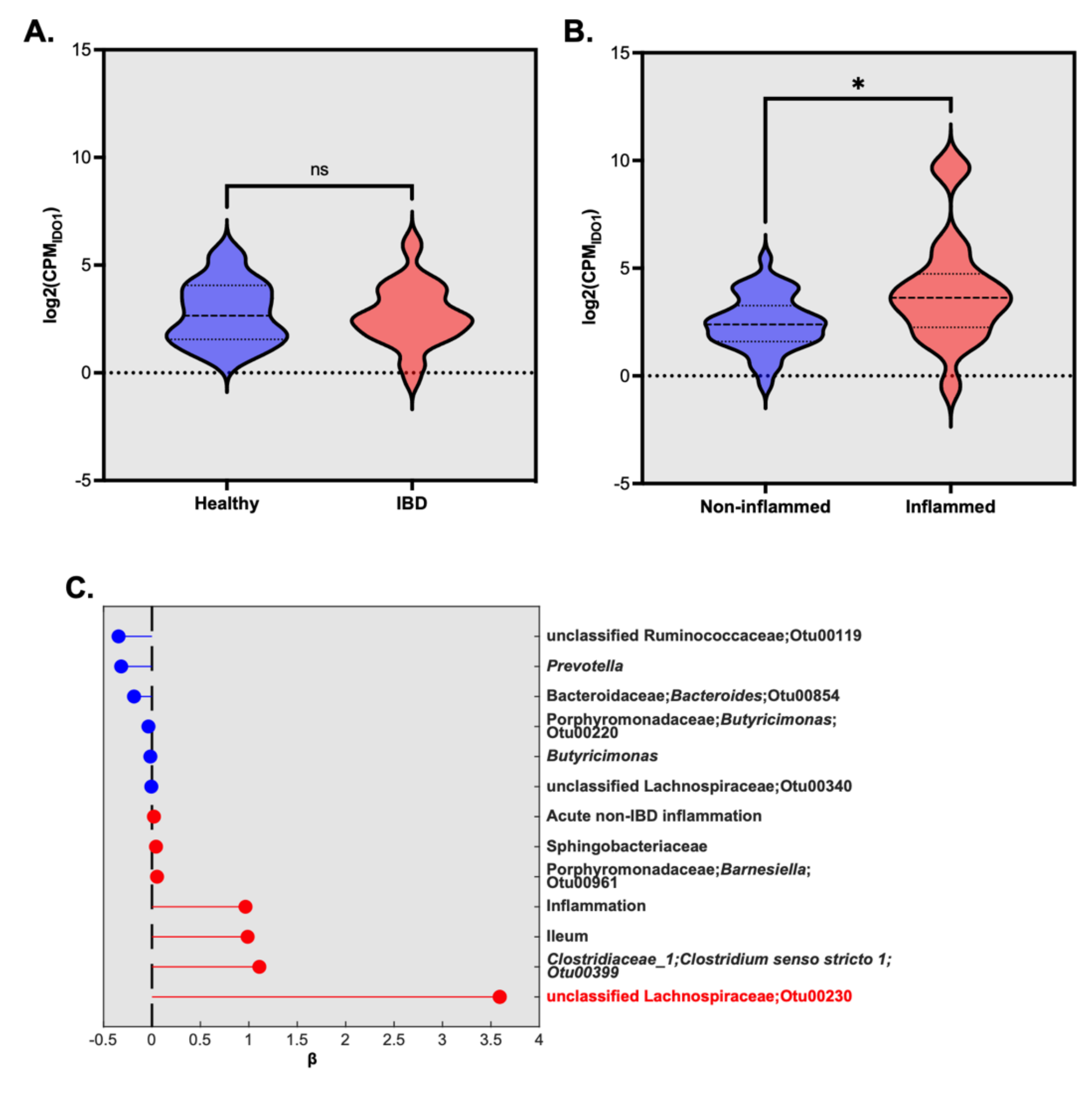


**Supplementary Figure 6. Characterization of IDO1 levels and identification of taxa that correlate with IDO1 levels in biopsies from the second IBD cohort re-analyzed in this study.** A) IDO1 levels do not differ between biopsies from healthy individuals and those with IBD. Ns- not significant, two-tailed Mann-Whitney test. B) IDO1 expression is significantly higher in biopsies collected from inflamed tissues compared to non-inflamed tissues. $P < 0.05$, two-tailed Mann-Whitney test. C) Taxa selected by LASSO regression as predictors of IDO1 levels in biopsies from this cohort. Red circles represent taxa positively correlated with IDO1, while blue circles represent taxa negatively correlated. Taxa highlighted in red belong to the Lachnospiraceae and Oscillospiraceae families. β: LASSO coefficient. ASV: amplicon sequence variant. CPM: counts per million. In violin plots, lines show median and quartile. Related to **Figure 6**.
